# HPV Oncogenes rewire DNA crosslink repair to create a PRIMPOL-dependent therapeutic vulnerability

**DOI:** 10.64898/2026.11.10.765685

**Authors:** Julien Dessapt, Marie-Anne Bérubé, Ottilie Le Doeuff, Nicolas Soucy, Line Balistaire, Béatrice Morin, Sadaf Valeh Sheida, Andréanne Blondeau, Yan Coulombe, Élise G. Lavoie, Jean-Yves Masson, Samer M. I. Hussein, Amélie Fradet-Turcotte

**Author notes:** Corresponding author : CHU de Québec - Université Laval Research Center (Hôpital de l’Enfant-Jésus) - Pavillon L 2260, boul. Henri-Bourassa Québec, QC, G1J 0J9. Contributed equally to the work (Julien Dessapt, Marie-Anne Bérubé).

## Abstract

Cisplatin-based chemoradiation is the standard of care for both HPV-negative (HPV^−^) and HPV-positive (HPV^+^) head and neck squamous cell carcinoma (HNSCC), even though patients with HPV^+^ tumors have better treatment responses and a better prognosis. Understanding how HPV status shapes this response could guide alternative strategies that lower long-term toxicity. Using genome-wide CRISPR screens, we show that the genetic requirements for surviving chemoradiation diverge between HPV^−^ and HPV^+^ HNSCC cells. HPV^+^ cells tolerate cisplatin-induced interstrand crosslinks independently of the Fanconi anemia pathway, a phenotype recapitulated by expressing HPV16 E6 and E7 in an HPV-negative background. In these cells, DNA damage tolerance depends instead on PRIMPOL-mediated repriming and post-replicative gap filling by translesion synthesis polymerases. Although they recruit BRCA1 normally to double-strand breaks, BRCA1 accumulation at cisplatin-stalled forks is markedly reduced, indicating that BRCA1 function is uncoupled between these two substrates. This tolerance pathway leaves cells dependent on dynamic RAD51 turnover at single-stranded DNA gaps. Depletion of the RAD51 unloading factor FIRRM, or chemical stabilization of RAD51 filaments, selectively kills HPV^+^ and HPV oncogene-expressing cells treated with cisplatin, and PRIMPOL depletion rescues this lethality. Our findings show that HPV oncogenes rewire the response to crosslink-induced replication stress and identify RAD51 filament dynamics as a therapeutic vulnerability in HPV^+^ HNSCC.

## INTRODUCTION

Human papillomaviruses (HPV) are small double-stranded DNA viruses that infect the keratinocytes of the skin and mucosa. Among the different types of HPV, high-risk mucosal alpha HPVs are responsible for about 4% of all cancers and are the etiologic agent of cervical cancers, a subset of anogenital and oropharyngeal cancers, the latter belonging to the Head and Neck Squamous Cell Carcinoma (HNSCC) group. HPV DNA is detected in an increasing proportion of oropharyngeal cancers. Clinically, the standard of care for both HPV-negative (HPV^-^) and HPV-positive (HPV^+^) HNSCC are the same, with concomitant radiotherapy and cisplatin-based chemotherapy^1^. Although having a positive HPV status in HNSCC is associated with a better prognosis^2^, these patients receive the same intensive treatment regimen as HPV^-^tumor, leaving long term survivors with significant acute and late toxicity including dysphagia, xerostomia, fibrosis, and feeding tube dependence^3,4^. While surgical de-intensification strategies rely on careful patient selection, non-surgical de-intensification trials have failed, compromising the survival of patients^5,6^. Understanding the molecular pathways that are activated by treatment in HPV^+^ vs HPV^-^tumors is key to developing alternative strategies for patients with HNSCC.

Cisplatin notably induces intrastrand and interstrand crosslinks (ICLs), the latter being highly cytotoxic DNA lesions acting as physical barriers to essential processes, including DNA replication, transcription, and recombination^7^. The detection, signaling and repair of these lesions involve a coordinated response triggered when the replication machinery encounters the ICL, which causes replication fork stalling and replication stress. The resolution of these lesions necessitates multiple pathways, including Fanconi anemia (FA), nucleotide excision repair (NER), and homologous recombination (HR) repair pathways, alongside the Ataxia Telangectasia and Rad3-Related (ATR) serine/threonine kinase checkpoint signaling pathways^7^. ICLs are recognized by the FANCM complex (FANCM, FAAP24, MHF1 and MHF2), which serves as a docking platform for seven additional FANC proteins (FANCA, B, C, E, F, G and L) together with FAAP20 and FAAP100, forming the multisubunit FA core complex. Acting as an E3 ubiquitin ligase with the E2 enzyme UBE2T/FANCT, this complex monoubiquitinates the FANCI-FANCD2 (ID2) heterodimer, the central step in ICL processing. Once activated, ID2 recruits the structure-specific nucleases that unhook the crosslink. It also serves as a hub for translesion synthesis polymerases, which bypass the residual adduct, and for the homologous recombination factors, which repair DNA double-strand breaks generated during unhooking^7^.

In addition to the FA pathway, ICLs can be bypassed by DNA damage tolerance (DDT) mechanisms, allowing replication to resume^8^. DDT includes translesion synthesis (TLS), in which specialized polymerases, including REV1, Polη, Polι, Polκ and Polζ, replicate across the damaged template, and template switching (TS), in which the nascent strand uses the sister chromatid as a template. A related mechanism, fork reversal, remodels the stalled fork into a four-way junction that repositions the lesion behind the fork, after which it can be repaired or bypassed by template switching. Alternatively, PRIMPOL-mediated repriming restarts synthesis downstream of the lesion, leaving a single-stranded DNA (ssDNA) gap between the adduct and the restarted fork^9^. This gap can then be filled post-replicatively by TLS or TS, allowing replication to continue with minimal effects on fork progression. The regulation and usage of these DDT mechanisms are emerging as a key factor influencing cancer therapy response across different subtypes of cancer^10,11^.

Here, we used genome-wide CRISPR-Cas9 knockout screens in HPV^-^ and HPV^+^ HNSCC cells to define the genetic requirements for surviving chronic chemoradiation and found that these requirements diverge sharply according to HPV status. HPV^+^ cells are sensitive to cisplatin, but in contrast to HPV^-^cells, those that survive do not depend on the FA pathway to tolerate cisplatin-induced lesions. Expression of HPV16 E6 and E7 is sufficient to recapitulate this phenotype, allowing us to attribute these observations to viral oncogenes rather than to differences in genetic background. In these cells, we show that survival depends on PRIMPOL-mediated repriming and post-replicative gap filling by TLS polymerases. Consistent with previous reports that HPV oncogene-expressing cells are HR-proficient^12,13^, these cells efficiently accumulated BRCA1 at DSBs. In contrast, BRCA1 accumulation at cisplatin-induced lesions is markedly reduced in HPV16 E6/E7-expressing cells, suggesting that these cells resolve ICLs either through a distinct mode of BRCA1 recruitment at stalled replication forks or independently of BRCA1. This tolerance program to DNA damage leaves E6/E7-expressing cells and HPV^+^ HNSCC cells dependent on dynamic RAD51 turnover at gaps. Under these conditions, stabilization of RAD51 filaments selectively kill HPV oncogene-expressing and HPV^+^ cells at cisplatin doses that spare their HPV^-^counterparts, in a PRIMPOL-dependent manner.

## RESULTS

### CRISPR-Cas9 knockout screens identify HPV status-dependent regulators of cisplatin and radiation sensitivity in HNSCC cells

To identify HPV status-dependent genetic vulnerabilities under chronic chemoradiation, we performed genome-wide CRISPR-Cas9 knockout screens in two oropharyngeal squamous cell carcinoma lines with different HPV status, UM-SCC-1 (SCC-1, HPV^-^) and UM-SCC-47 (SCC-47, HPV^+^)^14^. HPV and p53 status were confirmed by RT-qPCR for the expression of the viral oncogenes *E6* and *E7*, the tumor suppressor *TP53* and the p53 transcriptional targets *BAX* and *CDKN1A* (**Supplementary Fig.1A, Table S1**). Single-cell clones expressing a FLAG-tagged Cas9, SCC-1 Cas9 C10 (SCC-1 C10) and SCC-47 Cas9 C1 (SCC-47 C1) were selected based on Cas9 expression levels and editing efficiency at a locus containing of the locus *FAM83G*^15^, 6 days post-selection (**Supplementary Fig.1B-C**). For each clone, we used a chronic treatment regimen of cisplatin (2.5 nM, refreshed every 3 days) combined with ionizing radiation (0.125 Gy every 3 days), doses were titrated such that cumulative killing over the screen reached ∼80% relative to untreated cells (hereafter cumulative LD80)^16^. This was reach after 15 in days SCC-47 C1 and 21 days in SCC-1 C10 (**Supplementary Fig. 1D**). The screens were carried out using the TKOv1 single guide RNA (sgRNA) library (91,320 guides targeting 17,232 genes; 6 guides per gene)^17^ at a coverage of 400 per sgRNA and screened in biological duplicate for at least 10 population doublings with the control groups left untreated. Next-generation sequencing (NGS) of integrated sgRNA sequences was done at T0 and T15/21. DrugZ analyses identified fewer significantly depleted genes (NormZ score ≤ -2.3) in SCC-47 C1 when compared to SCC-1 C10. This was expected as the treated arm completed with fewer divisions in the HPV^+^ cell line. Nonetheless, core essential genes were significantly depleted in both cell lines with 52% in SCC-47 C1 and 63% in SCC-1 C10 (**Supplementary Fig. 1E**), confirming that both screens recovered known fitness genes with high specificity.

**Figure 1.**
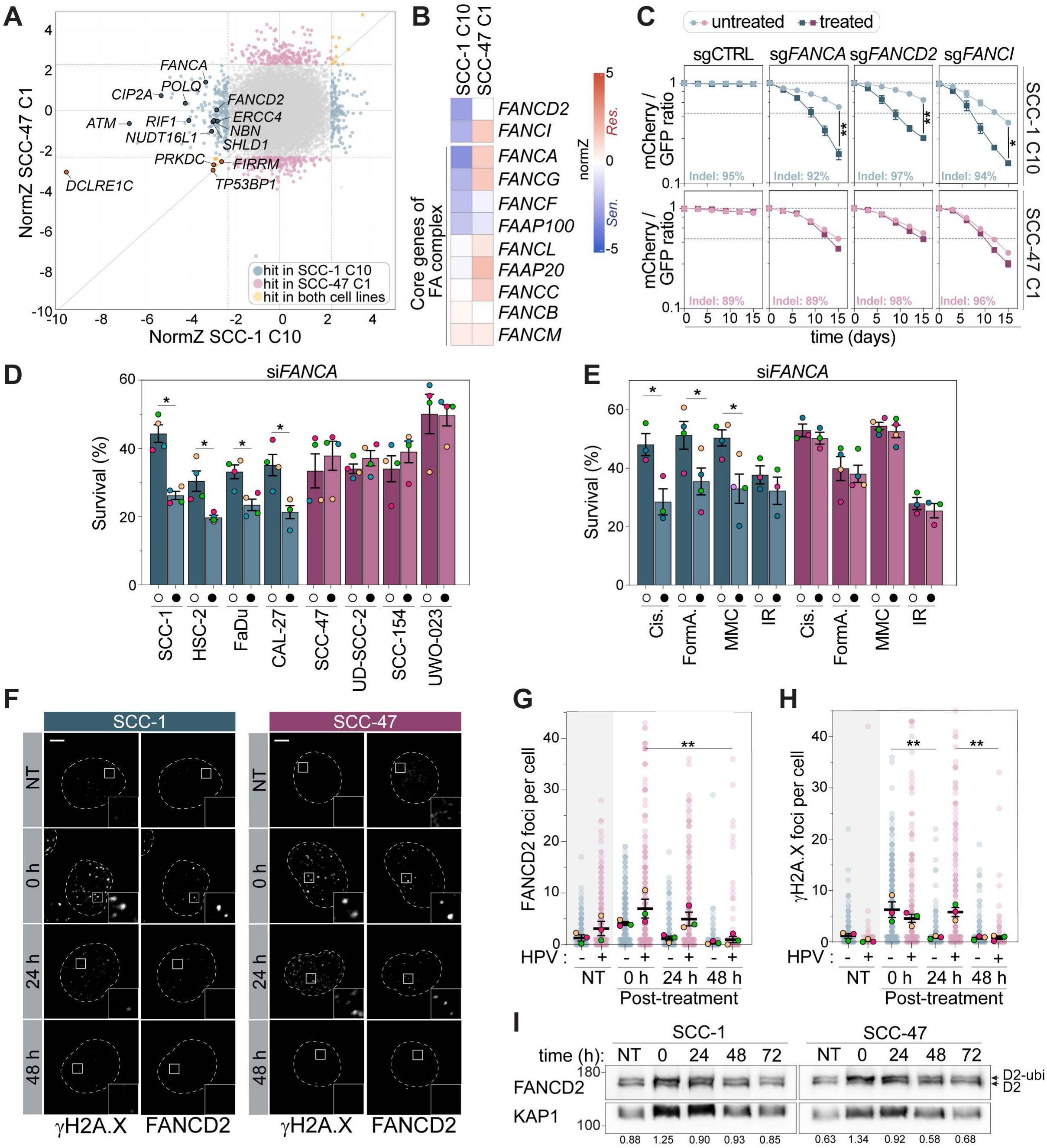
The Fanconi anemia pathway is engaged but dispensable for survival of chemoradiation in HPV+ HNSCC cells. **(A)** Pairwise CRISPR screens comparison performed in HPV^-^SCC-1 C10 and HPV^+^ SCC-47 C1 cells under chronic chemoradiation (IR 0.125 Gy and cisplatin 2.5 nM). Negative scores indicate sensitization and positive scores indicate resistance. Hits NormZ score ≤ -2.3 or ≥ 2.3 specific to SCC-1 C10 (blue), to SCC-47 C1(pink), or shared by both (orange). Selected DNA repair genes are labeled. **(B)** NormZ scores for core Fanconi anemia (FA) pathway genes in the screens shown in (A). **(C)** Competitive growth assays in Cas9-expressing SCC-1 C10 (blue) and SCC-47 C1 (pink) mixing GFP-labelled sg*FAM83G* (sgCTRL), or mCherry-labelled sg*FANCA*, sg*FANCD2* or sg*FANCI* cells. The mCherry/GFP ratio was monitored over 15 days without treatment (light) or under chronic chemoradiation (IR 1 Gy and cisplatin 20 nM; dark). Indel frequencies are given in each panel. **(D)** Survival of HPV^-^(blue) and HPV^+^ (pink) cell lines transfected with control (open circle) or *FANCA* (filled circle) siRNA and exposed to IR (12.5 Gy) and cisplatin (2 µM, 48 h). Open circles: CTRL siRNA; filled circles: indicated siRNA. **(E)** As in (D), for SCC-1 and SCC-47 cells treated with cisplatin (Cis.; 3 µM), formaldehyde (FormA.; 100 µM), mitomycin C (MMC; 75 nM) or ionizing radiation (IR; 12.5 Gy). **(F)** Representative γH2A.X and FANCD2 foci in SCC-1 and SCC-47 cells, untreated (NT) or fixed 0, 24 or 48 h after cisplatin treatment (250 nM, 24 h). **(G-H)** Quantification of (G) FANCD2 and (H) γH2A.X foci per cell in SCC-1 and SCC-47 cells, treated as in F. **(I)** Western blot of FANCD2 and monoubiquitinated-FANCD2 (D2-ubi) at the indicated times after cisplatin (2 µM, 24 h). KAP1 was used as a loading control. Numbers give the D2-ubi/D2 ratio. Data in (C–E) and (G–H) are mean of four independent biological replicates ± SEM; in (D, E, G, H), colored dots indicate the mean of each replicate and faded dots individual cells. * p ≤ 0.05 and ** p ≤ 0.01. Scale bars, 5 µm. See also Figure S1.

Genome-wide comparison of the two cell lines revealed reproducible depletion of distinct gene sets in the HPV^+^ vs HPV^-^background (**Fig. 1A**). Applying a NormZ score threshold of ≤ -2.3), 230 genes in SCC-1 C10 and 188 genes in SCC-47 C1 were significantly depleted. Among those, only 13 genes were shared between the two cell lines, including two core NHEJ factors (*DCLRE1C*, *PRKDC*) and *TP53BP1* (**Fig. 1A, Supplementary Fig. 1F**). Five additional components of NHEJ and the RIF1-shieldin end-protection axis (*RIF1*, *SHLD1*, *NUDT16L1*, *NHEJ1*) were significantly depleted in SCC-1 C10, validating the capacity of our screen to recover genes regulating DSB repair. Strikingly, *C1orf112/FIRRM* was robustly depleted in both cell lines, whereas FA pathway genes were selectively required in SCC-1 C10 (**Fig. 1A-B, Supplementary Fig. 1F**). FIRRM acts with FIGNL1 to promote RAD51 filament disassembly and the resolution of homologous recombination intermediates arising at ICLs^18–20^, suggesting that this function remains essential in the HPV^+^ cell line but is less dependent on the FA pathway.

To validate the specificity of the contribution of the FA pathway to survival in cells exposed to combined treatment with cisplatin and IR, we targeted the three most depleted FA genes (*FANCA, FANCD2* and *FANCI*) in a multicolor competitive growth assay^21^. Cell lines stably expressing a control sgRNA targeting *FAM83G* together with an NLS-GFP marker were mixed 1:1 with cells expressing an sgRNA against the gene of interest together with an NLS-mCherry marker, and the mCherry:GFP ratio was monitored every three days over 15 days of chronic cisplatin and IR treatment (**Fig. 1C, Supplementary Fig. 1G and Table S1**). Under untreated conditions, sgRNAs targeting *FANCA*, *FANCD2* and *FANCI* slightly reduced the relative fitness of both cell lines compared with sgCTRL (**Fig. 1C)**, consistent with additional functions of these factors in the protection of replication fork progression^22,23^.

Because our screen was performed in non-isogenic cell lines, we investigated whether this phenotype extended to a broader panel of HPV^-^ and HPV^+^ HNSCC cell lines (**Fig. 1D, Supplementary Fig. 1A**)^14,24,25^. Cells were transfected with an siRNA targeting *FANCA* or a control siRNA (siCTRL), for 24 h (**Supplementary Fig. 1H**) prior to treatment with irradiation and cisplatin. Survival was monitored 48 h post-treatment. Under these conditions, all cell lines treated with siCTRL showed between 30-50% survival. In contrast, the HPV^-^cell lines (SCC-1, HSC-2, FaDu, CAL-27), but not the HPV^+^ cell lines (SCC-47, UD-SCC-2, SCC-154, UWO-023) were further sensitized by the depletion of *FANCA* (**Fig. 1D**). We next used this assay to compare the sensitivity of SCC-1 and SCC-47 to different types of ICL-inducing agents (cisplatin, formaldehyde and mitomycin C)^7^ and to irradiation (**Fig. 1E**). As expected, the depletion of *FANCA* reduced the viability of SCC-1 cells treated with the ICL-inducing agents, but not that of irradiated cells. Consistent with our earlier findings, the viability of SCC-47 was unaffected by *FANCA* depletion under all conditions tested. Altogether, these results demonstrate that the FA-independent tolerance of HPV^+^ cells is neither cell line nor ICL agent specific. Importantly, *FANCA* depletion failed to sensitize either cell line to irradiation alone, indicating that the FA requirement observed under combined treatment is driven by the ICL-inducing component rather than by IR. The radioresistant origin of SCC-47 is therefore unlikely to account for its lack of FA dependency^26^.

The ability of HPV^+^ cells to survive ICLs without requiring the canonical FA pathway does not reveal whether the pathway is engaged or not, or whether it is activated but fails to progress. To distinguish between these possibilities, we monitored the formation and resolution of FANCD2 nuclear foci, together with γH2A.X levels as a readout of DNA damage (**Fig. 1F, H**). SCC-1 and SCC-47 cells were exposed to 250 nM cisplatin for 24 h and fixed either at the end of treatment (0 h) or 24 or 48 h after release into fresh medium. Both cell lines accumulated FANCD2 and γH2A.X foci with similar kinetics at 0 h, indicating that the FA pathway is engaged in HPV^+^ cells (**Fig. 1F, H**). However, resolution of these foci occurred later in SCC-47 than in SCC-1 (48 h vs 24 h, respectively), consistent with HPV^+^ cells processing ICL-induced damage through a distinct mechanism (**Fig. 1F-H**). In contrast, FANCD2 monoubiquitination peaked at the end of treatment and was resolved with comparable kinetics in both cell lines when monitored by western blotting analyses (**Fig. 1I**). We conclude that divergence between HPV^+^ and HPV^-^cells lies downstream of ID2 complex activation rather than in its activation.

### HNSCC cells expressing the viral oncogenes accumulate ssDNA

In keratinocytes, expression of HPV oncogenes E6 and E7 are essential for the development of HPV^+^ tumors. Expression of these oncogenes was previously reported to delay FANCD2 deubiquitination and impair ICL repair in a p53-dependent manner^27^. Although the HNSCC HPV^+^ cell lines used in this study are all *TP53* wild-type^14,25^, they are derived from tumors driven by two types of high-risk HPV (HPV16-driven: SCC-47, UD-SCC-2, SCC-154, and HPV33-driven: UWO023), isolated from different head and neck subsites and from patient at different stage of the disease^14,25,28^. To limit the contribution of genetic background difference, we engineered the *TP53*-mutant HPV^-^SCC-1 line^29^ to express both HPV16 E6 and E7 (E6E7). The retroviral vector pLXSN-16E6E7, in which E6 and E7 are cloned in their native viral arrangement (**Fig. 2A**, HPV16 nt 56-875, p97 through the E7 stop, E6 intron retained)^30–32^ and expressed from the Moloney murine leukemia virus promoter-enhancer located in the LTR of the retroviral vector was used to establish the cell line (SCC-1 16E6E7). A cell line transduced with the empty vector (EV) was generated as control. RT-qPCR showed that E6 and E7 transcripts accumulate to much lower levels in SCC-1 16E6E7 than in SCC-47 (**Supplementary Fig. 2A**). This is expected as SCC-1 16E6E7 was generated at a MOI of 0.3, which favors a single integration per cell, whereas six integration sites and 21-47 viral genome copies per cell have been mapped in SCC-47^33,34^. Despite these lower levels, E6/E7 expression was sufficient to recapitulate the phenotypes of the HPV^+^ HNSCC lines. SCC1-16E6E7 cells are more sensitive to cisplatin treatment than the SCC-1 EV, but surviving cells cope with cisplatin in a FANCA-independent manner (**Fig. 2B, Supplementary Fig. 2B**). In contrast, cell survival drops from 57 to 37% upon FANCA depletion in SCC-1 EV. In both cell lines, the resolution of FANCD2 foci (mean of 1.1 ± 0.48 foci in SCC-1 EV, 4.01 ± 1.1 foci in SCC-1 16E6E7 24 h post treatment) and γH2A.X foci (mean of 1.22 ± 0.36 foci in SCC-1 EV, 6.15 ± 0.35 foci in SCC-1 16E6E7 24 h post treatment) was delayed following treatment with cisplatin (**Fig. 2C-D, Supplementary Fig. 2C-E**). In contrast to the HNSCC HPV^+^ lines, but in line with previous work in keratinocytes^27^, FANCD2 deubiquitination was also delayed (**Supplementary Fig. 2F**). These findings indicate that the localization and deubiquitination of FANCD2 is independent of p53 status in these cells.

**Figure 2.**
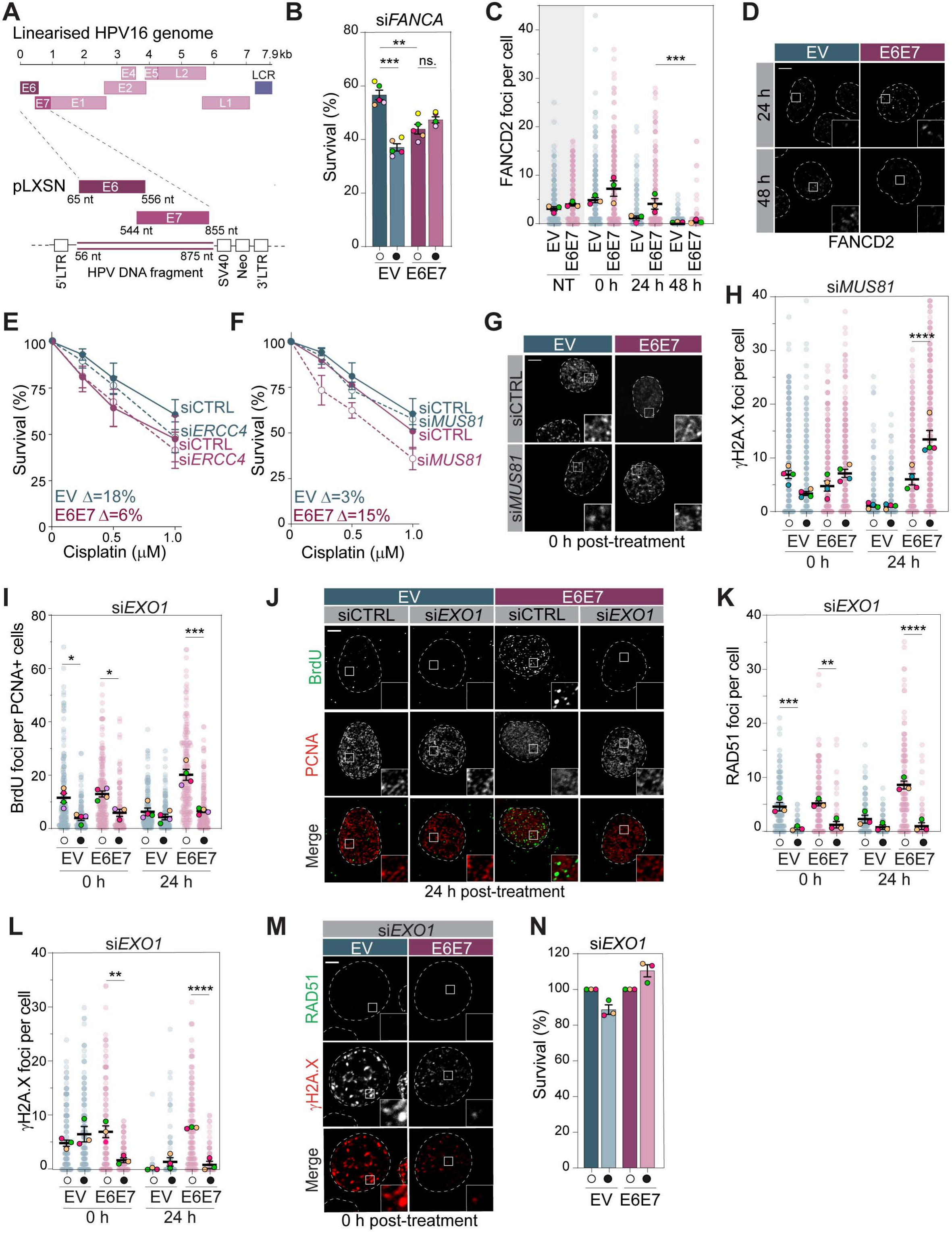
Cells expressing HPV16 E6 and E7 accumulate EXO1-dependent ssDNA after cisplatin. **(A)** Linearized HPV16 genome and the pLXSN retroviral construct used to express E6 and E7 (16E6E7). The HPV DNA fragment (nt 56-875), including the E6 (nt 65-556) and E7 (nt 544-855) open reading frames, is cloned between the 5′ and 3′ LTRs. SV40, SV40 promoter; Neo, neomycin resistance cassette. **(B)** Survival of SCC-1 cells stably transduced with pLXSN-16E6E7 or the empty pLXSN (EV), were transfected with control or *FANCA* siRNA and exposed to cisplatin (1 µM, 48 h). **(C, D)** Quantification (C) and representative images (D) of FANCD2 foci in SCC-1 EV and 16E6E7 cells, untreated (NT) or fixed 0, 24 or 48 h after cisplatin (250 nM, 24 h). Images show the 24 and 48 h time points. **(E, F)** Clonogenic survival of SCC-1 EV and 16E6E7 cells transfected with control, (E) *ERCC4/XPF*, or (F) *MUS81* siRNA and exposed to cisplatin (0-1 µM, 24 h). Δ, difference in survival between siCTRL and the indicated siRNA at 1 µM. **(G, H)** Representative images (G) and quantification (H) of γH2A.X foci in EV and 16E6E7 cells transfected with control or *MUS81* siRNA, fixed 0 or 24 h after cisplatin (250 nM, 24 h). Images show 0 h. **(I, J)** Quantification (I) and representative images (J) of BrdU foci per PCNA-positive cell in EV and 16E6E7 cells transfected with control or *EXO1* siRNA, fixed 0 or 24 h after cisplatin (2 µM, 24 h). Images show 24 h. **(K-M)** Quantification of RAD51 (K) and γH2A.X (L) foci per cell, and representative images (M), in cells transfected with control or *EXO1* siRNA, fixed 0 or 24 h after cisplatin (250 nM, 24 h). Images show the si*EXO1* condition at 0 h. **(N)** Survival of EV and 16E6E7 cells transfected with control or *EXO1* siRNA and exposed to cisplatin (1 µM, 48 h), normalized to the corresponding siCTRL (100%). Open circles, CTRL siRNA; filled circles, indicated siRNA. Faded dots, individual cells; colored dots, mean values of at least three biological replicates ± SEM. * p ≤ 0.05, ** p ≤ 0.01, *** p ≤ 0.001 and **** p ≤ 0.0001; ns, not significant. Scale bars, 5 µm. See also Figure S2.

Following the activation of ID2, unhooking of the cross-link then relies on nucleolytic incision by *ERCC4*/XPF-ERCC1, a process coordinated by the scaffold protein SLX4^7^. In our screen, the depletion of *ERCC4*/XPF is detrimental in SCC-1 (NormZ = -2.78) but not in SCC-47 (Norm Z -0.51) (**Fig. 1A, Supplementary Fig. 1F**), suggesting that HPV^+^ cells cope with cisplatin-induced ICLs without XPF-dependent unhooking. To test this, we measured the ability of *ERCC4*/XPF-depleted SCC-1 EV and 16E6E7 cells to recover from a 24 h exposure to 1 µM cisplatin using clonogenic assays. Consistent with the screen, *ERCC4*/XPF depletion reduced the survival of cisplatin-treated SCC-1 EV cells by 18%, compared with only 6% in 16E6E7 cells (**Fig. 2E, Supplementary Fig. 2G-H**). In contrast, MUS81 depletion reduced the survival of cisplatin-treated SCC-1 16E6E7 cells by 15%, while having little effect in EV cells (3%) (**Fig. 2F, Supplementary Fig. 2I-J**). Although MUS81 is not the main nuclease associated with ICL processing^7^, its depletion reduces γH2A.X levels in these cells after cisplatin treatment suggesting that it has at least a partial role in this condition (0 h post-treatment, **Fig. 2G-H, Supplementary Fig. 2K-L**). In contrast, the depletion of MUS81 in 16E6E7 increases the levels of γH2A.X, a phenotype that is even more striking 24 h after treatment (55.8% of 16E6E7 cells with more than 10 foci versus 2% of EV) (**Fig. 2G-H, Supplementary Fig. 2K-L**). MUS81 is a structure-specific endonuclease that associates with the non-catalytic subunit EME2 to cleave ICLs and promote fork restart in the S phase, and with EME1 to resolve recombination intermediates in G2/M^35^. The progressive accumulation of γH2A.X in MUS81-depleted 16E6E7 cells therefore suggests that these cells accumulate intermediates requiring MUS81-dependent resolution rather than MUS81-dependent unhooking.

To further define this phenotype, we investigated the levels of ssDNA that accumulate in both cell lines following cisplatin treatment. Cells were labelled with the nucleoside analogue BrdU for 24 h concurrently with cisplatin treatment, and ssDNA was detected with an anti-BrdU antibody under non-denaturing conditions (native BrdU)^36^. SCC-1 EV accumulated ssDNA after treatment, and this signal declined 24 h post-release, consistent with the lesion being repaired. In contrast, ssDNA in SCC-1 16E6E7 cells continued to increase over the same interval (**Fig. 2I-J, Supplementary Fig. 2M-O**), suggesting that the molecular events leading to ssDNA differs in the two cell lines. Nonetheless, ssDNA accumulation was dependent on the 5’-3’ exonuclease EXO1 in both cell lines as BrdU signal is abolished by its depletion. Similarly, we found the recombinase RAD51 specifically accumulates over time in 16E6E7 (**Fig. 2K, Supplementary Fig. 2P-Q**). Depletion of EXO1 in EV slightly increases the levels of γH2A.X after cisplatin treatment (**Fig. 2L-M, Supplementary Fig. 2Q-R**), consistent with DNA break being induced by XPF-ERCC1 unhooking prior to the accumulation of ssDNA. In contrast, depletion of EXO1 in 16E6E7 almost completely abolished γH2A.X signal, suggesting that ssDNA accumulation is causing DNA damage in this condition. This phenotype is reminiscent of conditions where γH2A.X accumulates following hyper resection by EXO1^37,38^. Although not significant in this context, EV cell survival is reduced upon depletion of EXO1 while it is increased in 16E6E7 (**Fig. 2N**), an observation that also support this model. Altogether, these results indicate that E6E7-expressing cells respond to cisplatin-induced ICLs through a pathway that requires XPF-ERCC1-dependent unhooking but involves EXO1-mediated resection followed by RAD51 loading on ssDNA. In this context, our results suggest that MUS81 contributes to the processing of DNA damage intermediates after ssDNA and RAD51 accumulation in E6/E7-expressing cells.

### PRIMPOL-dependent DNA gaps accumulated in 16E6E7 expressing cells following exposure to cisplatin

Beside the FA pathway, DNA damage tolerance mechanisms restore DNA synthesis by bypassing the obstacle rather than removing it^9,39^. One of these pathways relies on PRIMPOL-dependent repriming and allows DNA synthesis to resume synthesis downstream of the lesion, leaving post-replicative ssDNA gaps that are filled by translesion synthesis or template switching (**Fig. 3A**)^40,41^.

**Figure 3.**
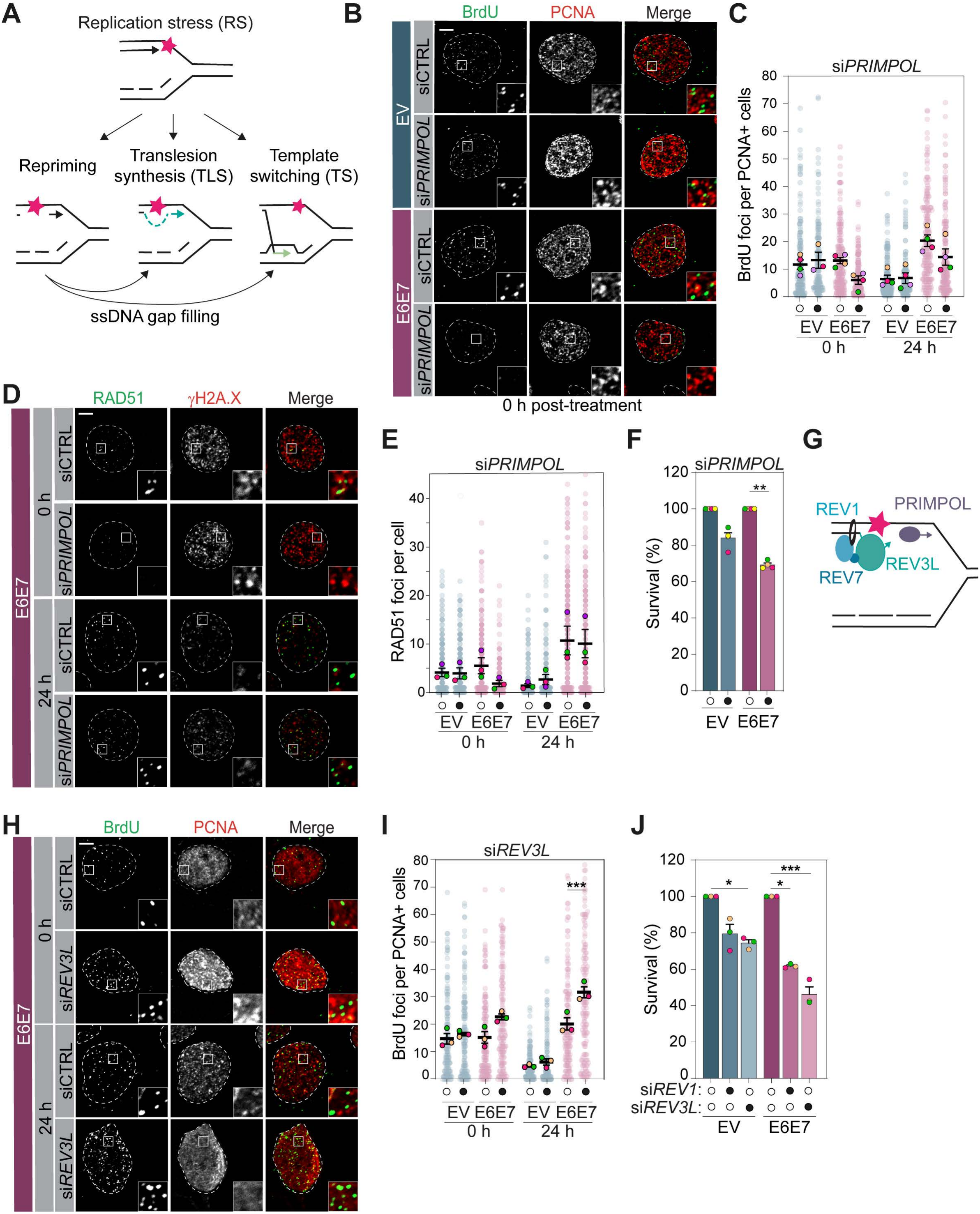
HPV16 E6E7-expressing cells accumulate PRIMPOL-dependent ssDNA gaps after cisplatin and rely on REV1/Rev3L-dependent TLS for survival. (**A**) Pathways processing a replication-blocking lesion. Repriming leaves ssDNA gaps behind the fork, which are filled by TLS or TS. (**B, C**) Representative images (B) and quantification (C) of BrdU foci per PCNA-positive cell in SCC-1 EV and E6E7 cells transfected with control (siCTRL) or *PRIMPOL* siRNA and fixed 0 or 24 h after cisplatin treatment (2 µM, 24 h). Images show 0 h. (**D, E**) Representative images of RAD51 and γH2A.X (D) and quantification of RAD51 foci (E) cells transfected as in (B) with cisplatin (250 nM, 24 h). Images show 16E6E7 cells at 0 and 24 h. **(F**) Survival of EV and 16E6E7 cells transfected with siCTRL or si*PRIMPOL* and exposed to cisplatin (1 µM, 48 h), normalized to the corresponding siCTRL (100%). (**G**) TLS complex (REV1, REV7, REV3L) and PRIMPOL at a stalled replication fork. (**H, I**) Representative images (H) and quantification (I) of BrdU foci per PCNA-positive cell in cells transfected with siCTRL or si*REV3L,* fixed 0 or 24 h after cisplatin (2 µM, 24 h). (**J**) Survival of EV and 16E6E7 cells transfected with siCTRL, si*REV1* or si*REV3L* and exposed to cisplatin (1 µM, 48 h), normalized to the corresponding siCTRL (100%). Open circles, CTRL siRNA; filled circles, indicated siRNA. Faded dots, individual cells; colored dots, mean values of at least three biological replicates ± SEM. * p ≤ 0.05, ** p ≤ 0.01 and *** p ≤ 0.001. Scale bars, 5 µm. See also Figure S3.

To test whether the observed accumulation of ssDNA in 16E6E7 cells is PRIMPOL-dependent, we quantified ssDNA (native BrdU foci) and RAD51 in control and PRIMPOL-depleted cells following cisplatin treatment. Both signals were reduced by about 2-fold upon PRIMPOL depletion in 16E6E7 but not in EV (**Fig. 3B-E, Supplementary Fig. 3A-E**). Despite this reduction in ssDNA, PRIMPOL-depletion was significantly detrimental to 16E6E7 cells but no to EV cells in survival assays (**Fig. 3F**). These findings indicate that repriming is not a by-product of damage but the DDT mechanism these cells depend on and suggest that fork arrests at ICL cannot be bypassed without PRIMPOL.

If repriming is productive only when the resulting gaps are subsequently filled, suppressing gap filling should be equally toxic. TLS polymerase σ (Pol σ), composed of the catalytic subunit REV3L and REV7, acts with the scaffolding protein REV1 to fill in the ssDNA gaps post-replicatively (**Fig. 3G**)^39^. Accordingly, depletion of REV3L increased the accumulation of ssDNA in 16E6E7 cells (**Fig. 3H-I, Supplementary Fig. 3F-H**) and sensitized them to cisplatin (**Fig. 3J**). Depleting REV1 produced a similar phenotype in survival assay (**Fig. 3J, Supplementary Fig. 3I**). These finding suggest that 16E6E7 cells rely on PRIMPOL-repriming followed by TLS-mediated gap filling to tolerate and interrupt any step that is lethal.

PRIMPOL-repriming protects stalled forks when fork reversal is inactivated^41^. This alternative strategy is particularly important for the tolerance of cisplatin-induced lesions in BRCA1-deficient cells^39–44^. HPV^+^ tumors, however, are not BRCA1 deficient as they do not display the SBS3 mutational signature associated with defective HR^13,45^ and cells expressing E6 and E7 repair DSBs efficiently by HR^12,13^. This raises the question of how PRIMPOL-dependent tolerance program is promoted in HPV^+^ cells. Strikingly, we found that, although BRCA1 foci accumulate efficiently at DSBs induced by IR (**Fig. 4A-B, Supplementary Fig. 4A, B**), BRCA1 foci accumulation at cisplatin-induced lesions was markedly reduced in 16E6E7 compared with EV cells (**Fig. 4C-D, Supplementary Fig. 4C**). Consistent with HR being required to repair DSBs generated by XPF-ERCC1/SLX4-dependent unhooking at ICLs^7^, depleting BRCA1 reduced both survival and RAD51 foci formation in EV but not in 16E6E7 cells (**Fig. 4E-G, Supplementary Fig. 4D-G**). Depleting CtIP also reduced RAD51 foci accumulation only in EV cells (**Fig. 4H-I, Supplementary Fig. 4H-I**). Although BRCA1 itself is dispensable at ICL in 16E6E7, RAD51 dynamics remain essential in both backgrounds. Depletion of the RAD51-loading factor BRCA2 abolished RAD51 foci formation in both cell lines (**Fig. 4J, Supplementary Fig. 4J-L**), and depletion of FIRRM, a component of the FIRRM/FIGNL1 RAD51 unloading complex, induced more cell death in 16E6E7 than in EV cells, specifically at low nM of si*FIRRM* (**Fig. 4K-L, Supplementary Fig. 4M**). These findings can be interpreted in two ways. First, impaired BRCA1 recruitment at cisplatin-induced lesions in 16E6E7 cells may favor PRIMPOL-dependent repriming, leading to the accumulation of ssDNA gaps and their resolution by post-replicative repair rather than by FA/HR-mediated ICL repair. Alternatively, reduced fork reversal in these cells may prevent the formation of the reversed-fork structures that BRCA1 is normally recruited to protect them against degradation by the MRE11 nuclease^22^ . These possibilities differ in whether reduced BRCA1 recruitment is a cause or a consequence of altered fork processing.

**Figure 4.**
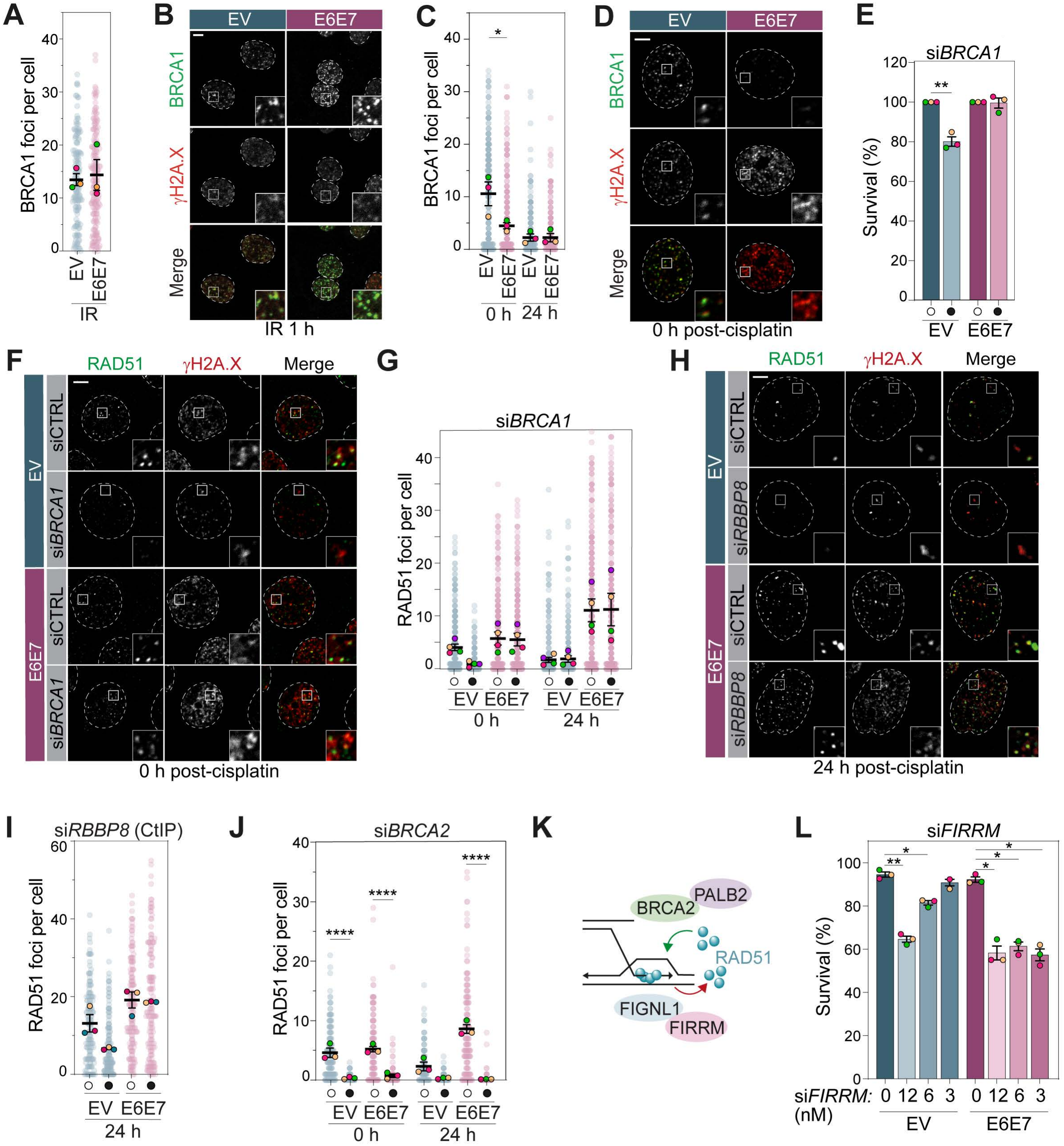
HPV16 E6E7-expressing cells form BRCA1-and CtIP-independent RAD51 foci. (**A, B**) Quantification (A) and representative images (B) of BRCA1 foci in SCC-1 EV and 16E6E7 cells fixed 1 h after ionizing radiation (IR; 0.5 Gy). (**C, D**) Quantification (C) and representative images (D) of BRCA1 foci in the same cells fixed 0 or 24 h after cisplatin (250 nM, 24 h). Images show 0 h. (**E**) Survival of EV and 16E6E7 cells transfected with control (siCTRL) or *BRCA1* siRNA and exposed to cisplatin (1 µM, 48 h), normalized to the corresponding siCTRL (100%). (**F, G**) Representative images (F) and quantification (G) of RAD51 foci in cells transfected with siCTRL or si*BRCA1*, fixed 0 or 24 h after cisplatin (250 nM, 24 h). Images show 0 h. (**H, I**) Representative images (H) and quantification (I) of RAD51 foci cells transfected with siCTRL or si*RBBP8*, fixed 24 h after cisplatin (2 µM, 24 h). (**J**) RAD51 foci per cell in cells transfected with siCTRL or si*BRCA2*, fixed 0 or 24 h after cisplatin treatment (250 nM, 24 h). (**K**) RAD51 regulation. BRCA2 and PALB2 promote RAD51 loading, FIGNL1 and FIRRM promote unloading. (**L**) Survival of EV and 16E6E7 cells transfected with si*FIRRM* (0, 3, 6 or 12 nM) and exposed to cisplatin (1 µM, 48 h). Open circles, CTRL siRNA; filled circles, indicated siRNA. Faded dots, individual cells; colored dots, mean values of at least three biological replicates ± SEM. * p ≤ 0.05, ** p ≤ 0.01 and **** p ≤ 0.0001. Scale bars, 5 µm. See also Figure S4.

### RAD51 dynamic is essential for the viability of 16E6E7 expressing cells

Our findings indicate that 16E6E7 cells depend more heavily on proper RAD51 turnover than on HR itself (**Fig. 4**), suggesting that disrupting the balance between RAD51 loading and unloading could represent a targetable vulnerability in HPV^+^ cancers. To test this hypothesis, we used small molecules that interfere with distinct steps of RAD51 filament assembly (**Fig. 5A**). RI-1 covalently modifies RAD51 at Cys319 and destabilizes the protein-protein interface required for filament formation^46^, while IBR2 disrupts RAD51 multimerization and promotes its proteasomal degradation^47^. Both molecules shift the equilibrium away from assembled filaments. In contrast, RS-1 allosterically locks RAD51 in an active conformation and stabilizes presynaptic filaments^32^. B02, which disrupts RAD51-binding to DNA and inhibits strand exchange^48,49^, was included as a fourth mechanistic class.

**Figure 5.**
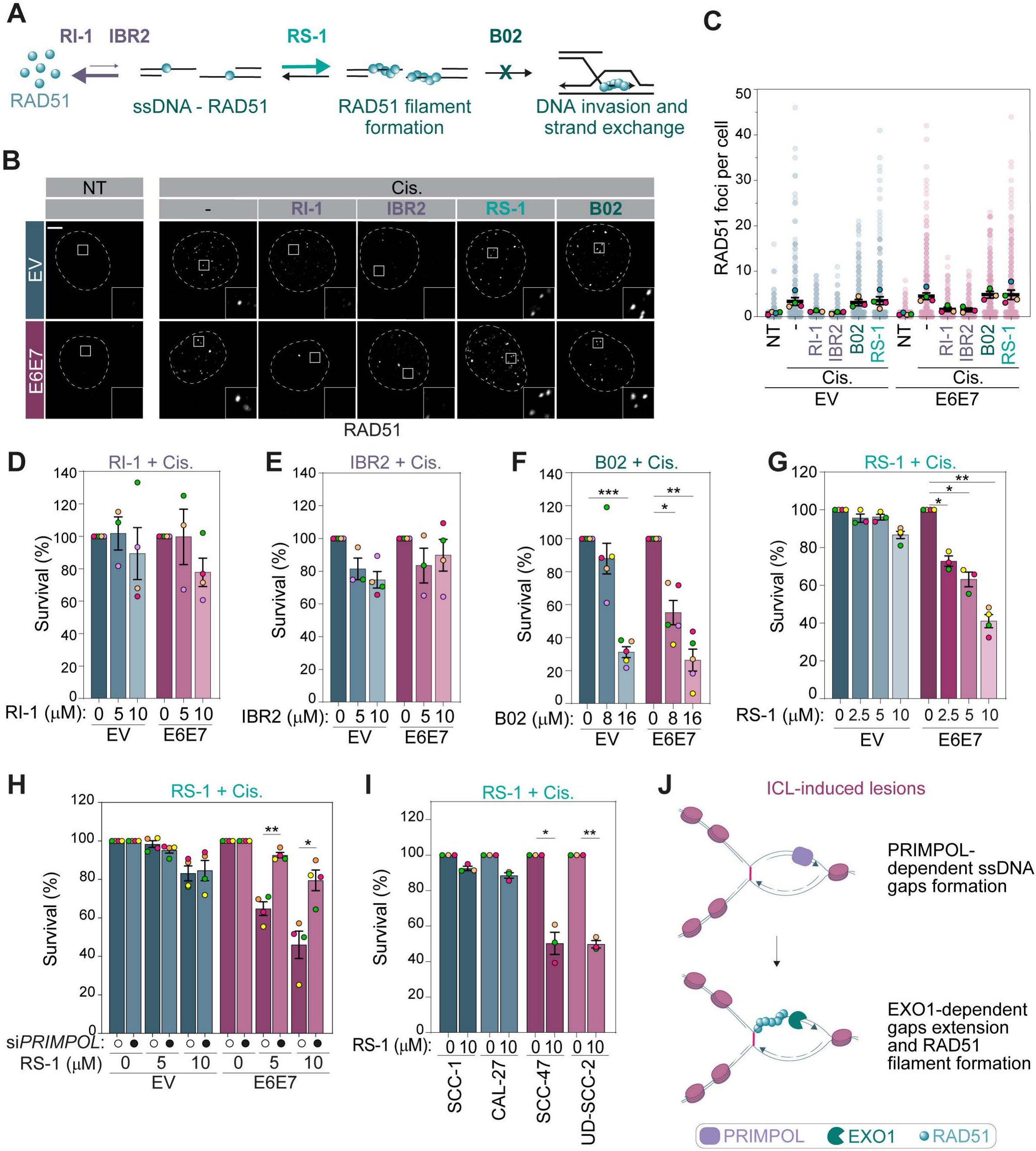
Stimulation of RAD51 activity selectively sensitizes HPV16 E6/E7-expressing and HPV-positive HNSCC cells to cisplatin in a PRIMPOL-dependent manner. (**A**) RAD51 loading and activation, and the step targeted of by each compound. RI-1 and IBR2 inhibit RAD51 filament formation on ssDNA, RS-1 stabilizes filament, and B02 blocks DNA invasion and strand exchange. (**B, C**) Representative images (B) and quantification (C) of RAD51 foci in SCC-1 EV and 16E6E7 cells, untreated (NT) or treated with cisplatin (250 nM, 24 h), alone or with RI-1 (10 µM), IBR2 (10 µM), B02 (8 µM) or RS-1 (5 µM). (**D-G**) Survival of EV and 16E6E7 cells exposed to cisplatin (500 nM, 48 h) with increasing concentrations of (D) RI-1 (0, 5 or 10 µM), (E) IBR2 (0, 5 or 10 µM), (F) B02 (0, 8 or 16 µM) or (G) RS-1 (0, 2.5, 5, 10 µM), normalized to the cisplatin-only condition (100%). (**H**) Survival of the same cells transfected with control or *PRIMPOL* siRNA and exposed to cisplatin (500 nM, 48 h) with RS-1 (0, 5 or 10 µM), normalized to the corresponding 0 µM RS-1 condition (100%). (**I**) Survival of HPV^-^(blue) and HPV^+^ (pink) cells exposed to cisplatin (500 nM, 48 h) or with RS-1 (10 µM), as in (D-G). (**J**) Model of the response of HPV16 E6/E7-expressing cells to ICL. ICL-induced replication stress is bypassed by PRIMPOL-dependent repriming downstream of the lesion, generating ssDNA gaps that are filled by TLS polymerases. Open circles, CTRL siRNA; filled circles, indicated siRNA. Faded dots, individual cells; colored dots, mean values of at least three biological replicates ± SEM. * p ≤ 0.05, ** p ≤ 0.01 and *** p ≤ 0.001. Scale bars, 5 µm. See also Figure S5.

When added concurrently with cisplatin, RI-1 (10 µM) and IBR2 (10 µM), reduced RAD51 foci by at least 50% in both cell lines, whereas neither RS-1 (5 µM) and B02 (8 µM) did (**Fig. 5B-C**). Consistent with the accumulation of RAD51 filament being toxic in cell treated with cisplatin^19^, only RS-1 and B02 reduced the viability of 16E6E7 cells at a cisplatin dose that was only slightly detrimental for cell survival (**Fig. 5D-G, Supplementary Fig. 5A**). The selectivity window was narrower for B02, which was toxic to both cell lines at the highest concentration tested, than for RS-1, which only modestly reduced EV viability at that dose (**Fig. 5F-G)**.

If this toxicity originates from the ssDNA gaps that accumulate in 16E6E7 cells, removing the source of those gaps should relieve it. Indeed, depletion of PRIMPOL significantly rescued the survival of RS-1-treated 16E6E7 cells (**Fig. 5H, Supplementary Fig. 5B),** directly linking PRIMPOL-dependent gap formation to the vulnerability to RAD51 filament stabilization. Together, these findings show that 16E6E7 cells rely on dynamic RAD51 turnover to survive cisplatin, and that this dependency constitutes a vulnerability that can be exploited to selectively kill HPV^+^ cells. Consistent with this hypothesis, RS-1 is only lethal in HPV^+^ cancer cells when tested on a subset of HPV^-^ and HPV^+^ cell lines (**Fig. 5I, Supplementary Fig. 5C)**.

## DISCUSSION

Locally advanced HPV^+^ and HPV^-^HNSCC are currently managed with a similar standard of care, relying on surgery and/or concurrent chemoradiation with cisplatin^50^, despite being biologically distinct diseases. HPV^+^ tumors inactivate p53 and Rb through viral oncoproteins rather than by mutations and carry a markedly better prognosis^1^. De-escalation has been proposed for HPV^+^ disease to reduce treatment-related toxicity, but uniform dose reduction has not matched standard chemoradiation in randomized testing (NRG-HN005), underscoring the need to better understand the molecular mechanisms engaged by these tumors in response to treatment before alternative strategies can be designed. Here, we show that HPV oncogene expression promotes PRIMPOL-dependent repriming and TLS-mediated gap filling at ICL-induced fork stalling, a pathway usually activated in HR-deficient cells or in cells where fork reversal is restrained^40,43^. In cells expressing the viral oncogenes, this DDT program uncouples gap-based cisplatin tolerance from HR status and leaves these cells dependent on dynamic RAD51 turnover. As this phenotype is specifically observed in cells expressing the viral oncogenes upon cisplatin treatment, targeting that dependency provides a conceptual basis for a tumor-intrinsic approach to sparing normal tissue, distinct from uniform de-escalation.

Among the factors required in both HPV^−^ and HPV^+^ screens, FIRRM/C1orf112 was the only one acting in the replication-associated response to ICLs, whereas the FA pathway was selectively required in HPV^-^cells. Because FIRRM acts with FIGNL1 as an anti-recombinase during ICL repair^18–20,51^, we propose that this shared requirement reflects the dependency of HPV^+^ cells on RAD51 turnover at PRIMPOL-dependent gaps. Consistent with this interpretation, we show that FIRRM depletion is more lethal in cells expressing HPV16 E6 and E7 than in control cells, and chemical stabilization of RAD51 filaments reproduced this selectivity. These findings argue that the shared depletion in our screens does not simply reflect a general fitness requirement for restraining RAD51 filament accumulation. Rather, HPV^+^ cells operate within a narrow tolerance for interference with RAD51 dynamics, such that even a small change in filament stability is sufficient to compromise survival when treated with cisplatin. This model predicts that direct inhibition of the FIRRM-FIGNL1 complex should phenocopy the selectivity we observe with RS-1. Small molecules targeting FIGNL1 have been recently described^52^; testing whether they selectively reduce the viability of HPV^+^ cells in parallel treatment with cisplatin could be an interesting avenue to test this prediction.

An important question left open by our data concerns the fate of the crosslink itself. We show that the FA core complex is recruited and that ID2 is monoubiquitinated with close to normal kinetics in HPV^+^ and 16E6E7 cells, yet XPF-ERCC1 is not required for survival. Whether a nuclease unhooks ICLs has not been tested. However, the depletion of MUS81 increases γH2A.X in 16E6E7 cells while it greatly reduces them in EV cells, suggesting that this nuclease does not act at the ICL unhooking step in 16E6E7 cells.

A related question concerns which cisplatin lesion drives gap formation. Intrastrand adducts vastly outnumber ICLs in cisplatin-treated cells^53^ and can be bypassed by repriming without any requirement for traverse or replisome remodelling, so the ssDNA gaps we measure may arise at either class of lesion. The fact that FA-independence of HPV^+^ cells extends to mitomycin C and formaldehyde argues that the divergence we describe is driven by the crosslinking activity of these agents rather than by intrastrand damage but formally establishing that the gaps themselves form at ICLs will require comparing gap induction by cisplatin with that of agents such as melphalan, which produced an higher proportion of intra-crosslinks^54^. An alternative route to ICL tolerance should also be considered that FANCM has been reported to promote replication traverse of ICLs^55,56^. The recruitment of FANCM and subsequent traverse depend on its translocase activity, FAAP24 and ATR, but are independent of the FA core complex and ID2. FANCD2 promotes, but is not required for, restart^55,56^. Whether FANCM-dependent traverse allows the replisome to bypass ICLs that persist in E6/E7 expressing cells cannot be distinguished by the assays used here. The latter possibility is attractive, as it would explain why ID2 activation proceeds normally while the downstream requirements diverge, and it predicts that crosslinks should persist across divisions in these cells. Direct measurement of ICL removal by modified comet assay or crosslinked-plasmid reactivation will be required to resolve this.

Replication stress refers broadly to replication fork challenges, but it is becoming increasingly clear that fork repair mechanisms are highly tailored toward specific types of damage and replication intermediates. How cells choose between fork reversal, repriming and translesion synthesis at a stalled fork, and how these pathways are regulated across the cell cycle, remains poorly defined^8^. Our data identify HPV oncogenes expression as one determinant of that choice. In cells expressing HPV16 E6 and E7, BRCA1 accumulation at cisplatin-induced lesions is markedly reduced, yet BRCA1 is recruited efficiently to DSBs induced by ionizing radiation in the same cells. This selectivity is unexpected given current models of BRCA1 recruitment to damaged DNA. At DSBs, BARD1, through its ARD-BRCT module, targets the BRCA1-BARD1 complex to damaged chromatin by reading unmethylated H4K20 on post-replicative chromatin together with RNF168-dependent H2AK15 ubiquitination^57–59^. The mechanism by which E6/E7 restricts BRCA1 at one substrate and not the other is unknown.

At ICL-induced lesions, BRCA1 contributes to the disassembly of the CMG helicase^60^ prior to nuclease degradation of the ICL and protects reversed forks from nucleolytic degradation by MRE11 following fork reversal^61,62^. Because CMG unloading is a prerequisite to the incision-dependent FA route^7^, restricting BRCA1 at ICL-stalled forks would be predicted to preserve an engaged replisome, the configuration compatible for PRIMPOL to reprime downstream of the lesion. Thus, it is unclear which complex of BRCA1 is detected at ICL-induced lesions and even whether these complexes are differently regulated. It is yet to be understood whether differential recruitment of BRCA1 is directly regulated by E6 and E7 or a consequence of the engagement of a different DDT pathway. Indeed, PRIMPOL-dependent ssDNA gaps and fork reversal are mutually exclusive replication stress tolerance pathways^9^. The PRIMPOL-dependent ssDNA gaps pathway is favored in cells where fork reversal is inhibited and in cells where the levels of PRIMPOL are increased^39,40,43,63^. Whether and how the viral oncogenes interfere with fork reversal or the level of PRIMPOL are unknown and distinguishing these possibilities will require direct assessment of fork reversal in these cells. Favouring repriming over reversal would provide a means of avoiding the pathological reversed-fork intermediates that arise when fork protection is compromised, a strategy that may also be advantageous during the productive viral life cycle in differentiating keratinocytes and retained as a by-product of oncogene expression in transformed cells.

Our experiments do not resolve which viral oncogene drives this phenotype, as E6 and E7 were expressed together throughout. One contribution can nonetheless be excluded: SCC-1 cells carry a *TP53* mutation, so the phenotype arises independently of E6-mediated p53 degradation. E7 is an attractive candidate, as we have previously shown that it interacts with RNF168^12^. RNF168-dependent ubiquitination governs the recruitment of BRCA1-BARD1 to damaged chromatin through the H2AK15ub mark and contributes to the recruitment of SLX4 at ICLs^64^. Interference with this single E3 ligase could therefore account for both the restricted BRCA1 accumulation and the altered requirements for SLX4-associated nucleases that we observe in E6/E7-expressing cells. Testing this will require expressing each oncogene separately and determining whether RNF168 activity is limiting for either event.

Altogether, our findings reveal that HPV^+^ cells fail to accumulate BRCA1 at cisplatin-induced lesions and adopt a tolerance program otherwise characteristic of BRCA1-deficient cells exposed to chronic cisplatin treatment^40,43^ . This occurs despite normal BRCA1 recruitment to DSBs, in line with previous reports that HPV^+^ cancer cells are HR-proficient^13^. These observations show that BRCA1 function can be uncoupled between DSBs and stalled replication forks. Identifying this difference raises the interesting possibility of exploiting gap-centric synthetic lethality. In this case, the prediction would be that HPV^+^ tumors are PARPi-sensitive despite being HR-proficient and that persistent ssDNA in HPV^+^ tumors predict ATR dependence. Beyond HPV, these findings illustrate that gap-based drug tolerance can arise in cells with intact HR, and that the genomic scars used to infer repair capacity may not predict which replication-associated vulnerabilities a tumor carries.

## METHODS

### EXPERIMENTAL MODEL AND SUBJECT DETAILS

#### Cell lines

All cell lines were maintained at 37°C in a humidified incubator with 5% CO₂. All culture media were supplemented with 10% v/v fetal bovine serum (FBS; 12483020; Thermo Fisher Scientific). UM-SCC-1 (male; SCC070; MilliporeSigma) and UM-SCC-47 (male; SCC071; MilliporeSigma) were maintained in Dulbecco’s Modified Eagle’s Medium / Nutrient Mixture F-12 (DMEM/F12) (11330057; Thermo Fisher Scientific). UD-SCC-2, FaDu, Cal-27, UWO23, HSC-2, and UPCI-SCC-154 (all male) were a kind gift from Anthony Nichols and John Barrett (Western University, London, Ontario, Canada) and were cultured in DMEM/F12. All HPV^+^ cell lines were of HPV16 subtype except for UWO23 which is HPV33. Expression of the HPV oncogenes E6 and E7 was confirmed by RT-qPCR in all cell lines. The cell lines were confirmed to be mycoplasma-free using a PCR mycoplasma detection kit (ABMG238; Diamed) and were routinely tested every six months.

## METHOD DETAILS

### Plasmids

Additional plasmid information is provided in the Key Resources Table. LentiCas9-Blast, used to generate the Cas9-expressing cell lines for the CRISPR screens, was a gift from Feng Zhang (Addgene plasmid #52962; RRID: Addgene_52962)^65^. The Toronto human knockout pooled library (TKO.v1) was a gift from Jason Moffat (Addgene #1000000069)^17^. For retrovirus production, pUMVC was a gift from Bob Weinberg (Addgene plasmid #8449; RRID: Addgene_8449). For lentivirus production, psPAX2 and pMD2.G were gifts from Didier Trono (Addgene plasmids #12260 and #12259; RRID: Addgene_12260 and RRID: Addgene_12259). For competitive growth assays, sgRNAs (Table S1) were cloned using BsmBI into LentiCRISPR-NLS-GFP-sgRNA (control sgRNA) or LentiCRISPR-NLS-mCherry-sgRNA (gene of interest), both gifts from Dr. Dan Durocher (The Lunenfeld-Tanenbaum Research Institute, Toronto)^21^. pLXSN16E6E7 was a gift from Denise Galloway (Addgene plasmid #52394; RRID: Addgene_52394)^31^.

### Cell line generation, culture, and treatment

Stable Cas9-expressing UM-SCC-1 and UM-SCC-47 cell lines were established using the LentiCas9-Blast vector. Briefly, lentiviruses were produced by co-transfecting the vector with envelope-expressing plasmid pMD2.G (#12259; Addgene) and packaging plasmid psPAX2 (#12260; Addgene) using polyethyleneimine hydrochloride (PEI Max; 24765; Polyscience Inc.). After 48 h of transfection, supernatants containing lentiviral particles were collected and used to transduce UM-SCC-1 and UM-SCC-47 cells in the presence of 8 µg/mL hexadimethrine bromide (Polybrene; TR-1003; MilliporeSigma) and 10 mM HEPES pH 7.6. After 2 days of transduction, infected cells were selected with 6 µg/mL blasticidin S hydrochloride (BLA477; BioShop). Single-cell clones were obtained by limiting dilution, screened for Cas9 expression by western blotting, and validated for Cas9 activity using an sgRNA targeting *FAM83G* (sgFAM83G). Stable Cas9-expressing UM-SCC-1 and UM-SCC-47 clonal lines were transduced with a lentiviral vector expressing sgFAM83G and selected with 1 µg/mL puromycin for 6 days. Genomic DNA was extracted (QIAGEN QIAamp DNA Blood Mini Kit, 51104), the target region was amplified by PCR, and the amplicons were digested with SacII (NEB, R0157S) and resolved on an agarose gel. Because the sgRNA target site overlaps a SacII site, Cas9-induced indels render the amplicon resistant to digestion, so editing was detected by an undigested band.

UM-SCC-1 HPV16 pLXSN cell lines were generated by retroviral transduction with pLXSN16 containing E6 and E7 (pLXSN16E6E7), and control UM-SCC-1 cells were generated with the empty pLXSN16 vector (EV). Retroviruses were produced similarly to lentiviruses using the envelope-expressing plasmid pMD2.G (#12259; Addgene) and the packaging plasmid pUMVC (#8449; Addgene). The cells were infected at a MOI of 0.3 and selected with 500 µg/mL G418 Sulfate (BS723; Biobasic) for 14 days, and the expression of HPV16 oncogenes was confirmed using RT-qPCR.

For experiments using siRNAs, unless stated otherwise, cells were transfected with a total of 12 nM of individual or pooled siRNAs (see Table S1) using Lipofectamine RNAiMAX Transfection Reagent (13778150; Thermo Fisher Scientific) according to the manufacturer’s protocol. After 24 h, the cells were either left untreated, treated with cisplatin (2251; Tocris), with formaldehyde (CAJT2106-1; J.T.Baker), with mitomycin C (M0503; Sigma-Aldrich), with B02 (HY-101462; Cedarlane), with RI-1 (HY-15317; Cedarlane), with IBR2 (HY-103710; Cedarlane), with RS-1 (S8234; Selleckchem), with VE-821 (SML1415; Sigma-Aldrich), or exposed to IR in a CellRad+ irradiator (Precision X-Ray). Dosages used for each treatment are specified for each experiment. Cells were processed 48 h after siRNA transfection to validate the efficacy using western blotting or RT-qPCR.

### CRISPR-Cas9 Based Genome-Wide Screening

UM-SCC-1 and UM-SCC-47 Cas9-expressing cells were exposed to the TKO v1 sgRNA lentiviral library (#1000000069; Addgene) at a multiplicity of infection (MOI) of 0.3, and by ensuring coverage of at least 400-fold for each individual sgRNA within the cell population. One day post-transduction, puromycin selection (2 μg/mL) was carried out for 6 days to enrich for successfully transduced cells. Following selection, cells were divided into two experimental pools at a density of 4.5 million cells per plate (50 plates total per screen; D0). Pool 1 was treated with chronic cisplatin (2.5 nM) and ionizing radiation (IR; 0.125 Gy) at a dose reducing cell number by 80% relative to untreated after 15 to 21 days (LD80), while Pool 2 received vehicle. Over 15-21 days of culture, cells were maintained with puromycin (1 μg/mL) and passed every 3 days at a density of 4.5 million cells/plate to sustain 400-fold coverage per sgRNA. Cell pellets from treated and untreated conditions were collected at each time point and frozen at -80°C. Genomic DNA was then extracted with the QiAamp Blood Maxi Kit (Qiagen, Cat# 51194).

### NGS Library Preparation and Sequencing

sgRNA sequences were amplified from genomic DNA using two rounds of nested PCR. Outer PCR was performed using Kapa HiFi Ready Mix polymerase (7958935001; Roche) with forward and reverse outer primers (Table S1). PCR conditions: 18 cycles, 66°C annealing temperature, 15 seconds extension time. PCR products were pooled, and 4% of the input was re-amplified to incorporate Illumina HiSeq adapter sequences (Table S1.1). Products were purified using the QIAquick gel extraction kit (Qiagen) following separation on a 2% agarose gel. Deep sequencing was performed on the HiSeq 4000 (Illumina) platform (∼25M reads per sample, single end read, 250x sequencing depth).

### CRISPR screen data analysis

Raw sequencing files (FASTQ) were assessed for quality using FastQC. The MAGeCK^66^ count command was used to determine the library adapter length, perform an initial alignment against the TKO_v1 library, and generate sgRNA read counts. Reads were then trimmed according to the adapter length and re-aligned to the TKO_v1 library with Bowtie2^67^ using standard parameters. Final read count tables for each condition were regenerated from the Bowtie2-aligned reads with MAGeCK count (v0.5.9.3). Changes in sgRNA abundance after treatment were analyzed with DrugZ (v1.1.0.2), an algorithm developed for chemogenetic CRISPR screens, using default parameters (paired samples, pseudocount of 5 added to each sgRNA count). DrugZ^68^ returned gene-level normalized Z-scores (NormZ), calculated from up to 6 sgRNAs per gene. Screen quality was assessed by comparing the dropout rate over time of core essential genes with that of core non-essential genes, as previously described. Sensitizer hits were defined as genes with a NormZ < −2.3 at the final timepoint.

### CRISPR-Cas9 Based Competition Growth Assay

Competition-based growth assays were performed following established protocols. Lentiviral particles expressing LentiCRISPR-NLS-GFP-sgRNA-FAM83G or LentiCRISPR-NLS-mCherry-sgRNA-GOI targeting FANCA, FANCD2, or FANCI were used to transduce UM-SCC-1 and UM-SCC-47 cells at a MOI of 0.3. sgRNA sequences are presented in Table S1. Following transduction, cells were subjected to puromycin selection (2 μg/mL) for 6 days. GFP- and mCherry-labeled cells were then combined at a 1:1 ratio and seeded in 12-well plates. Beginning 1 day post-seeding (T1) and every 3 days thereafter, cells were treated with IR (1 Gy) and Cisplatin (20 nM; LD80 dose) or vehicle control. Fluorescence intensity of each population was measured by flow cytometry using a BD Accuri C6 Plus Flow Cytometer System (BD Biosciences). Cells were maintained and subcultured for a total of 15 days, with the mCherry-to-GFP fluorescent cell ratio determined by flow cytometry every 3 days and was analyzed using FlowJo software v10.10 (BD Biosciences).

### Cell viability and clonogenic cell survival assays

Cells were seeded in 6-well plates 24 h before transfection with the indicated siRNA. After 24 h, cells were harvested, counted, and divided from the same transfected pool for use in two parallel assays. For clonogenic cell survival assays, transfected cells were reseeded at 1,000 cells/well in new 6-well plates and treated 24 h later according to the predetermined cisplatin gradient. The medium was replaced with fresh medium 24 h after treatment, and cells were incubated for a further 12 days. Colonies were fixed in 100% methanol for 10 min at -20°C and stained with 0.1% (v/v) crystal violet in deionized water for 10 min at room temperature. Wells were rinsed with water and air-dried. The dye was then solubilized in 20% (v/v) acetic acid, and absorbance at 590 nm was measured using a BioTek Cytation 5 multimode reader with Gen5 software (Agilent). For short term viability assays, cells were seeded in new 6-well plates. After 24 h, cells were exposed to the indicated compound for 48 h. Cells were then counted manually under the microscope with an hemacytometer, and viability was calculated as the ratio of treated to vehicle-treated samples for each condition.

### Cellular extract preparation and immunoblotting

Cells were lysed either directly in 2× Laemmli buffer (500 mM Tris-HCl pH 6.8, 6% w/v SDS, 60% v/v glycerol, 0.03% w/v bromophenol blue, and 60 mM DTT) or in high-salt buffer (300 mM NaCl, 50 mM Tris-HCl pH 7.5, 1 mM EDTA, and 1% v/v Triton X-100; TB0198; BioShop) supplemented with 1× complete EDTA-free Protease Inhibitor Cocktail (11873580001; MilliporeSigma), 1 mM NaF, and 0.2 mM Na₃VO₄. High-salt lysates were subsequently denatured by heating in 6× Laemmli buffer for 10 min at 95^°^C. Protein concentrations were determined using the Pierce™ BCA Assay Kit (PI23227; Thermo Fisher Scientific). Equal amounts of protein were resolved by SDS-PAGE, transferred to nitrocellulose (10600002; Cytiva) or PVDF membranes (IPVH00010; Cytiva), and probed with the indicated primary antibodies. HRP-conjugated secondary antibodies were used for chemiluminescent detection with Radiance ECL (AC2204; Azure Biosystems). For BRCA1 and BRCA2 western blot, cells were lysed in a specific buffer (50 mM Tris-HCl pH 7.5, 0.1% w/v SDS, 20 mM NaCl, 1mM MgCl2) supplemented with 1× complete EDTA-free Protease Inhibitor Cocktail and benzonase (706643; Thermo Fisher Scientific). Lysates were subsequently denatured by heating in 6× Laemmli buffer for 2 min at 95^°^C, loaded on a NuPAGE Novex 3-8% Tris-Acetate gel (WG1603A; Thermo Fisher Scientific), and probed with the indicated primary and secondary antibodies.

### Antibodies

The following antibodies were used for western blotting: anti-FANCA (A301-980A-M; Invitrogen), anti-FANCD2 (NB100-182; Novus), anti-FANCI (sc-271316; Santa Cruz Biotechnology), anti-KAP1 (A300-274A; Cedarlane), anti-XPF (13465; Cell Signaling Technology), anti-REV1 (sc-393022; Santa Cruz Biotechnology), anti-EXO1 (ab95068; Abcam), anti-RAP80 (ab124763; Abcam), anti-glyceraldehyde 3-phosphate dehydrogenase (GAPDH; 39-8600; Thermo Fisher Scientific), anti-SMARCAL1 (sc-376377; Santa Cruz Biotechnology), anti-HLTF (ab183042; Abcam), anti-a-Tubulin (CP06; MilliporeSigma), anti-a-Actin-1 (A2066; MilliporeSigma), anti-Mus81 (NB100-2064; Novus), anti-FIRRM (ab121774; Abcam), anti-CtIP (61142; Active Motif), anti-FIGNL1 (HPA055542; MilliporeSigma), anti-BRCA1 (07-43; MilliporeSigma), anti-BRCA2 (OP95; MilliporeSigma). Secondary antibodies used were goat anti-Rabbit IgG, horseradish peroxidase (HRP; 111-035-144; Jackson ImmunoResearch Laboratories Inc.) and rabbit anti-mouse-HRP (A9044; MilliporeSigma). Primary antibodies used for immunofluorescence were: anti-FANCD2 (NB100-182; Novus), anti-RAD51 (70-001, BioAcademia), anti-phospho-Histone H2A.X (Ser139; 05-636; MilliporeSigma), anti-phospho-Histone H2A.X (Ser139; 2577L; Cell Signaling Technology), anti-BrdU (ab6326; Abcam), anti-BrdU (347580; BD Biosciences), anti-PCNA (NBP2-67390, Novus), anti-BRCA1 (07-434; MilliporeSigma). Secondary antibodies for immunofluorescence were from Thermo Fisher Scientific: Goat anti-Rabbit IgG, Alexa Fluor 488 (A-11034), 555 (A-21428), and 647 (A-21244); goat anti-Mouse IgG, Alexa Fluor 488 (A-11029), 555 (A-21424), and 647 (A-21236).

### RNA extraction and RT-qPCR

Total RNA was isolated using the RNeasy Mini Kit (74104; QIAGEN Inc.) or the Monarch® Spin RNA Isolation Kit Mini (T2110L; New England Biolabs®) according to the manufacturer’s instructions. Genomic DNA was digested on-column using the New England Biolabs kit and by DNase I digestion (1 unit; EN0521; Thermo Fisher Scientific) at 37°C for 2 h using the QIAGEN kit. Reverse transcription was then performed with 500 ng of RNA using the High-Capacity cDNA Reverse Transcription Kit (4368814; Thermo Fisher Scientific), following the manufacturer’s specifications. Quantitative PCR amplification was performed on a QuantStudio 3 Real-Time PCR System (Thermo Fisher Scientific) with LightCycler 480 SYBR Green I Master Mix (04707516001; MilliporeSigma), with 40 cycles of 95°C for 15 s, 56°C for 15 s and 72°C for 15 s, or with Power SYBR™ Green PCR Master Mix (4367659; Thermo Fisher Scientific), with 40 cycles of 95°C for 15 s, 56°C for 15 s, and 60°C for 1 min, using 1% of the cDNA reaction as the template. Gene expression quantification was performed via standard curve analysis using GAPDH, VAPA, RSP13 or PUM1 for normalization. Primer sequences are presented in Table S1.

### Immunostaining and imaging

UM-SCC-1 (WT and pLXSN-E6E7) and UM-SCC-47 cells were plated on coverslips in 24-well culture dishes. After exposure to the specified treatments and/or transfections, samples were subjected to fixation following one of two protocols: (1) incubation in 2% w/v paraformaldehyde (P6148; MilliporeSigma) in PBS for 20 min at ambient temperature, or (2) immersion in 100% methanol for 20 min at -20°C, the latter being used exclusively for RAD51 detection. Paraformaldehyde-treated samples underwent three rounds of PBS (270 mM NaCl, 5.4 mM KCl, 15.5 mM Na₂HPO₄ (SPD307; BioShop), 3 mM KH₂PO₄ (PPM302; BioShop), pH 7.4) rinses, followed by permeabilization with 0.3% v/v Triton X-100 in PBS for 20 min at ambient temperature. Cells were then incubated for 30 min in one of two blocking solutions: 10% v/v goat serum (16210064; Thermo Fisher Scientific), 0.5% v/v NP-40, and 0.5% w/v saponin (S7900; MilliporeSigma) in PBS, or alternatively, 2% w/v BSA. Primary antibody incubation proceeded for 2 h at ambient temperature. Three PBS washes preceded a 1 h incubation with secondary antibodies conjugated to Alexa Fluor dyes and DAPI counterstain (0.8 µg/mL; D3571; Thermo Fisher Scientific). Coverslips were then mounted onto slides using ProLong Diamond Antifade Mountant (P36970; Thermo Fisher Scientific) following final PBS washes. Imaging was performed on a ZEISS LSM 900 confocal microscope using 40× and 63× oil objectives. Quantitative analysis relied on ImageJ (National Institutes of Health) processing with a tailored macro routine: nuclear segmentation via binary masking, foci identification by "Find Maxima" with experiment-specific brightness/contrast parameters, and threshold calibration based on vehicle-treated reference samples. Dashed lines indicate nuclear boundaries in figures lacking DAPI visualization, and inset magnifications are 20×.

### BrdU ssDNA detection assay

siRNA-transfected cells were treated with 2 μM cisplatin in the presence of 10 μM BrdU (B23151; Thermo Fisher Scientific) for 24 h. Cells then underwent sequential chromatin extraction using a nuclear pre-extraction buffer (10 mM PIPES pH 7.0, 100 mM NaCl, 300 mM sucrose, 3 mM MgCl₂, 1 mM EGTA, 0.5% Triton X-100) followed by a cytoskeleton stripping buffer (10 mM Tris pH 7.4, 10 mM NaCl, 3 mM MgCl₂, 1% Tween-20, 0.5% sodium deoxycholate). Samples were rinsed three times with PBS and fixed sequentially: first with 2% w/v paraformaldehyde for 20 min on ice, then with 100% methanol for 5 min at -20°C. Permeabilization was conducted with 0.5% Triton X-100 for 10 min on ice, followed by blocking with 3% w/v BSA in PBS for 1 h on ice. Cells were then incubated overnight at 4°C with primary antibodies against BrdU and PCNA. Following three PBS washes, secondary incubation was performed with anti-mouse Alexa Fluor 488 and anti-rabbit Alexa Fluor 555 for 1 h at room temperature, succeeded by DAPI counterstaining (0.4 μg/mL) for 5 min at room temperature. Coverslips were mounted and imaged as previously described.

### Quantification and Statistical Analysis

All experiments were performed in at least triplicate biological replicates, except where specified otherwise. Quantification was performed on a minimum of 100 nuclei per condition per replicate, except for BrdU ssDNA detection assays, in which 50 PCNA-positive nuclei were analyzed per replicate. Data presentation incorporated SuperPlots methodology as outlined by Lord et al.^69^ to simultaneously display individual biological replicates and aggregated statistics. Graphical depictions show mean values ± standard error of the mean (SEM). Multi-group datasets displayed as SuperPlots were analyzed by nested one-way ANOVA followed by Tukey’s post hoc test for pairwise comparisons. Other multiple comparisons were performed by one-way ANOVA followed by Tukey’s multiple comparisons test. Two-group comparisons were performed using unpaired t-tests with Welch’s correction. All tests were performed on non-normalized data using GraphPad Prism v9.5.0 (GraphPad Software, San Diego, CA, USA). Significance thresholds were defined as * p ≤ 0.05, ** p ≤ 0.01, *** p ≤ 0.001, **** p ≤ 0.0001; ns, not significant.

## RESOURCE AVAILABILITY

### Lead Contact

Amelie Fradet-Turcotte

## ACKNOWLEDGMENTS

We thank members of the Fradet-Turcotte laboratory for reading and commenting on the manuscript and Roger Greenberg for the U2OS 2-6-5 cell line. We thank C. St-Pierre from the imaging unit of the CHU de Québec-Université Laval Research Center for his assistance in microscopic analyses. A.F.T. was supported by the J.-Louis Lévesque Foundation. A.F.T, S.M.I.H. and J.Y.M are supported by Canadian Institutes of Health Research grant (PJT-470053, PJT-378019, and PJT-173246, respectively). M.A.B and O.L.D are supported by doctoral fellowships from the Cancer Research Society (CRS) and Fonds de Recherche du Québec -Santé (FRQ-S), respectively. N.S. is supported by a master fellowship from FRQ-S. A.F.T. and J.Y.M hold Canada Research Chairs in Molecular Virology and Genomic Instability (Tier II), and DNA repair and Cancer Therapeutics (Tier I). Work in A.F.T lab was supported by the Fonds de Recherche du Québec through the research center grant for the CHU de Québec-Université Laval Research Center (30641).

## AUTHOR CONTRIBUTIONS

A.F.T. conceived and funded the project, designed the research, and wrote the manuscript. J.D. and M.A.B designed, performed, and analyzed most experiments, wrote the first draft and reviewed the manuscript. O.L.D, L.B., B.M., N.S., S.V.F, A.B., Y. C., and E.G.L. performed and analyzed RNA extraction, RT-qPCR, control immunoblotting experiment and flow cytometry. O.L.D., N.S., and A.B. also reviewed the manuscript. J.Y.M., and S.M.I.H. provided resources, expertise, and manuscript feedback.

