## Supplementary material for "HPV Oncogenes rewire DNA crosslink repair to create a PRIMPOL-dependent therapeutic vulnerability": Supplemetal figures 1-5 and Table 1

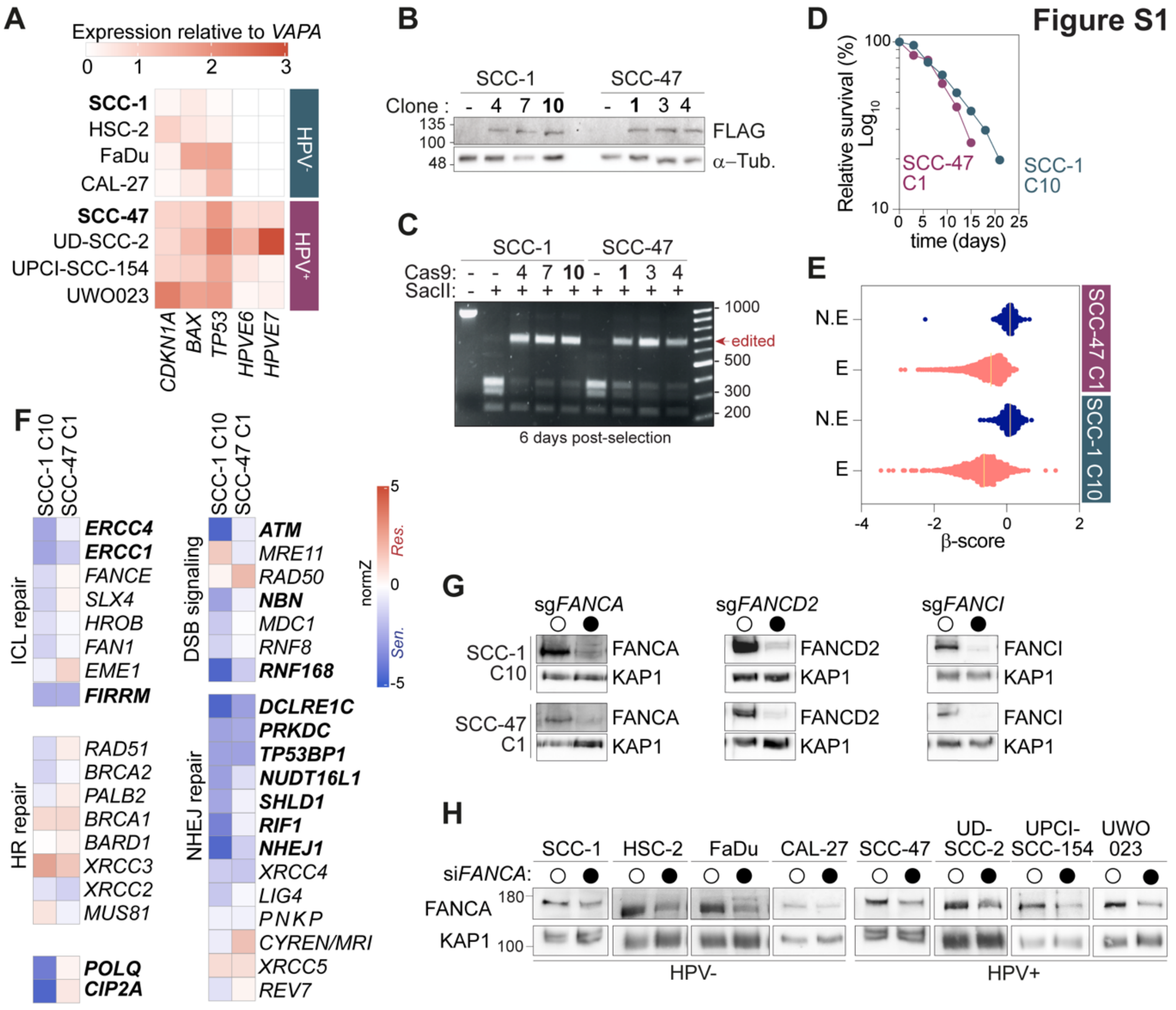


**Figure S1. Characterization of the HPV- and HPV+ HNSCC models and validation of the screen hits**

**(A)** Heatmap of *CDKN1A, BAX, TP53, HPV E6* and *HPV E7* expression measured by RT-qPCR in HPV- and HPV+ HNSCC cell lines, relative to *VAPA*. Cell lines used for the screens are in bold. **(B)** Western blot of FLAG-Cas9 in SCC-1 and SCC-47 clones. -, parental cells. Tubulin was used as a loading control. Clones SCC-1 C10 and SCC-47 C1 (bold) were selected for the screens. **(C)** Cas9 activity in the indicated clones 6 days post-selection. The *FAM83G* locus was PCR-amplified and digested with SacII. Editing disrupts the SacII site, giving an undigested product (edited). **(D)** Relative survival of SCC-1 C10 and SCC-47 C1 cells under chronic chemoradiation (0.125 Gy and 2.5 nM cisplatin). **(E)** Distribution of β-scores for essential (E) and non-essential (N.E) genes in the SCC-1 C10 and SCC-47 C1 screens. **(F)** Clustered NormZ scores for genes acting in ICL repair, HR repair, DSB signaling and NHEJ repair. Significant hits in either cell line are in bold. **(G)** Western blot validating FANCA, FANCD2 and FANCI depletion in SCC-1 C10 and SCC-47 C1 cells expressing sgCTRL (open circles) or the indicated sgRNA (filled circles). KAP1 was used as a loading control. **(H)** Western blot validating FANCA depletion by siRNA in the indicated cell lines. KAP1 was used as a loading control. See also Figure 1.

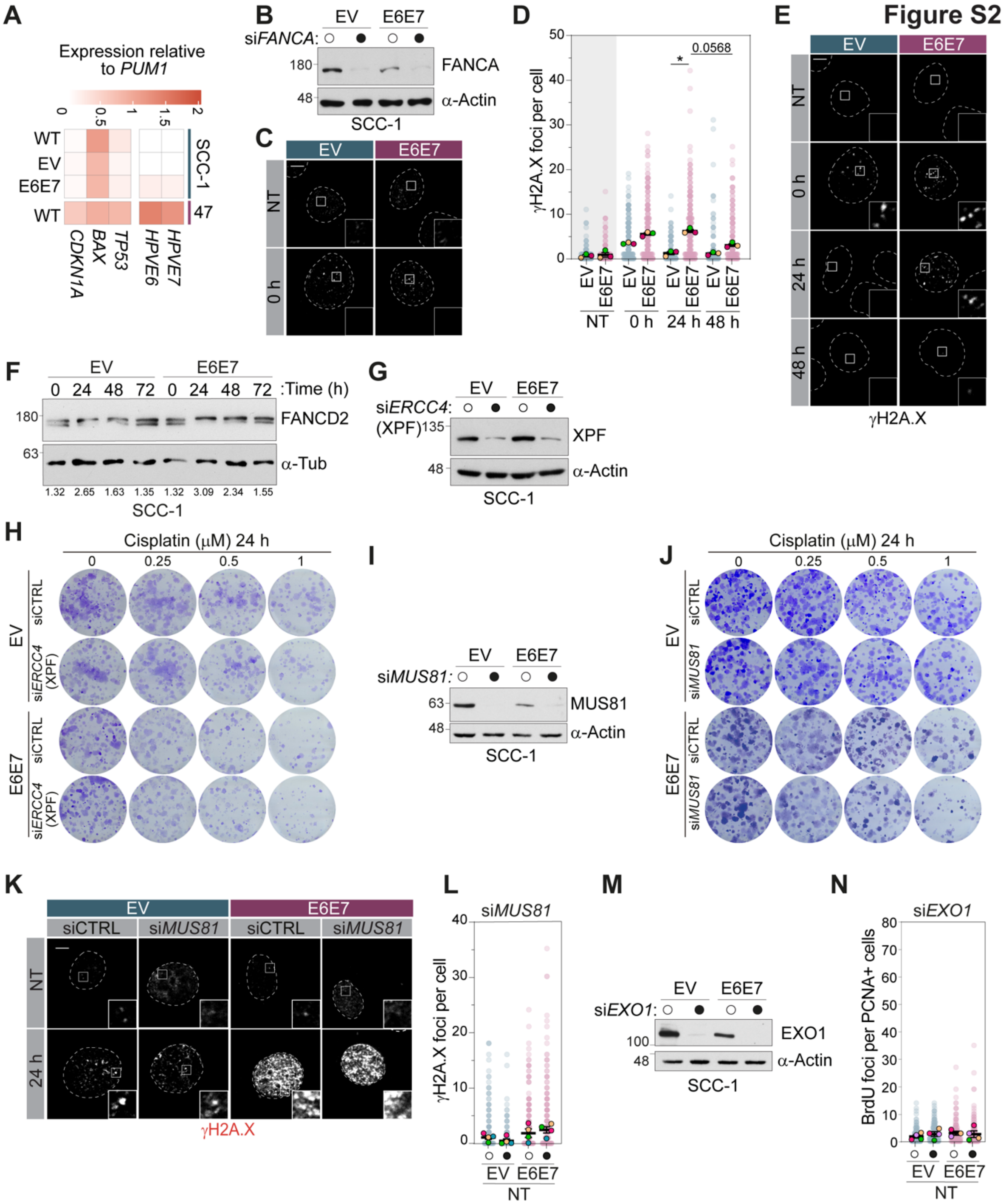
Figure S2
A
Expression relative to PUM1
0 0.5 1 1.5 2
WT
EV
E6E7
WT
SCC-1
47
CDKN1A
BAX
TP53
HPVE6
HPVE7
B
EV
E6E7
siFANCA:
180
FANCA
48
α-Actin
SCC-1
C
EV
E6E7
NT
0 h
D
γH2A.X foci per cell
0.0568
EV E6E7 EV E6E7 EV E6E7 EV E6E7
NT 0 h 24 h 48 h
E
EV
E6E7
NT
0 h
24 h
48 h
γH2A.X
F
EV
E6E7
0 24 48 72 0 24 48 72 :Time (h)
180
FANCD2
63
α-Tub
1.32 2.65 1.63 1.35 1.32 3.09 2.34 1.55
SCC-1
G
EV
E6E7
siERCC4:
(XPF)
135
XPF
48
α-Actin
SCC-1
H
Cisplatin (μM) 24 h
0 0.25 0.5 1
EV
E6E7
siCTRL
siERCC4 (XPF)
I
EV
E6E7
siMUS81:
63
MUS81
48
α-Actin
SCC-1
J
Cisplatin (μM) 24 h
0 0.25 0.5 1
EV
E6E7
siCTRL
siMUS81
K
EV
E6E7
siCTRL
siMUS81
NT
24 h
γH2A.X
L
siMUS81
γH2A.X foci per cell
EV E6E7
NT
M
EV
E6E7
siEXO1:
100
EXO1
48
α-Actin
SCC-1
N
siEXO1
BrdU foci per PCNA+ cells
EV E6E7
NT

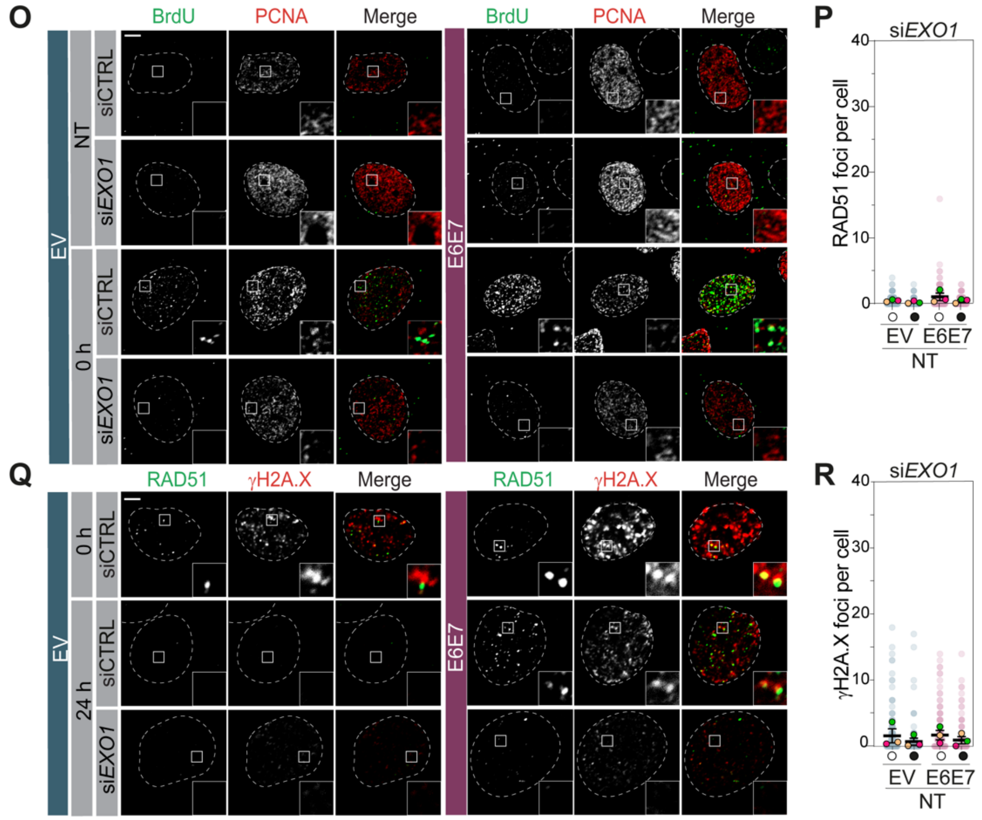


**Figure S2. Characterization of the SCC-1 16E6E7 model and validation of siRNA depletions** **(A)** Heatmap of *CDKN1A, BAX, TP53, HPV E6* and *HPV E7* expression measured by RT-qPCR in parental (WT), empty vector (EV) and E6E7-expressing SCC-1 cells, relative to *PUM1*. Parental HPV$^{+}$ SCC-47 cells are shown for comparison. **(B, G, I, M)** Western blot validating FANCA (B), XPF (si*ERCC4*) (G), MUS81 (I), EXO1 (M) depletion in SCC-1 EV and 16E6E7 cells. α-Actin was used as a loading control. **(C)** Representative images of FANCD2 foci in SCC-1 EV and 16E6E7 cells, untreated (NT) or fixed 0 h after cisplatin (250 nM, 24 h). **(D, E)** Quantification (D) and representative images (E) of γH2A.X foci per cell in the same cells, untreated (NT) or fixed 0, 24 or 48 h after cisplatin (250 nM, 24 h). **(F)** Western blot of FANCD2 and its monoubiquitinated form at the indicated times after cisplatin (1 µM, 24 h). α-Tubulin was used as a loading control. Numbers give the D2-ubi/D2 ratio. **(H, J)** Representative colony formation assays quantified in Figure 2E and 2F, in cells transfected with siCTRL or *ERCC4* (H) or si*MUS81* (J) exposed to the indicated cisplatin concentration for 24 h. **(K, L)** Representative images (K) and quantification (L) of γH2A.X foci in cells transfected with siCTRL or si*MUS81*, untreated (NT) or fixed 24 h after cisplatin (250 nM, 24 h). (L) shows untreated cells. **(N, O)** Quantification (N) and representative images (O) of BrdU foci per PCNA-positive cell in cells transfected with siCTRL or si*EXO1*. (N) shows untreated cells (NT), (O) untreated cells and fixed 0 h after cisplatin (2 µM, 24 h). **(P, Q, R)** Quantification of RAD51 (P) or γH2A.X (R) foci and representative images of RAD51 and γH2A.X (Q) in cells transfected with siCTRL or si*EXO1*. (P) shows untreated cells, (Q) 0 and 24 h post-cisplatin and (R) untreated cells. Treated conditions are shown in Figure 2. Open circles, CTRL siRNA; filled circles, indicated siRNA. Faded dots, individual cells; colored dots, mean values of at least three biological replicates ± SEM. * $p \leq 0.05$, an exact p-value is given where close to significant. Scale bars, 5 µm. See also Figure 2.

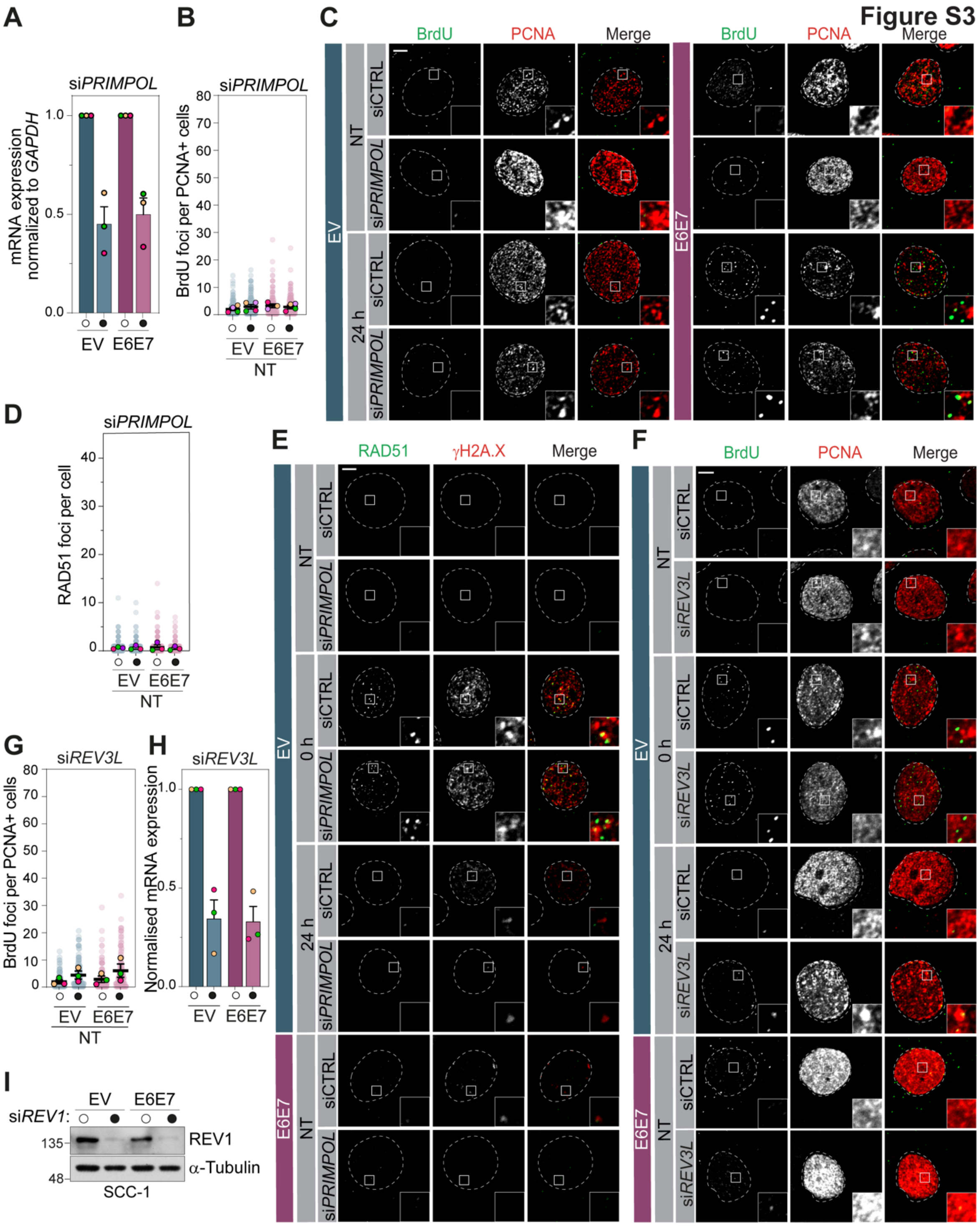


**Figure S3. Validation of PRIMPOL, REV3L and REV1 depletions and supporting data on ssDNA gaps.** (**A, H**) RT-qPCR validating *PRIMPOL* (A) and *REV3L* (H) depletion in SCC-1 EV and 16E6E7 cells, normalized to *GAPDH* and expressed relative to the corresponding siCTRL. (**B, C**) Quantification (B) and representative images (C) of BrdU foci per PCNA-positive cell in cells transfected with control (siCTRL) or *PRIMPOL* siRNA. (B) shows untreated cells (NT) and (C)

shows untreated cells and cells fixed 24 h after cisplatin (2 μM, 24 h). (**D, E**) Quantification (D) and representative images (E) of RAD51 foci in cells transfected with siCTRL or si*PRIMPOL*. (D) shows untreated cells and (E) shows EV cells left untreated or fixed 0 or 24 h after cisplatin (250 nM, 24 h), and 16E6E7 cells untreated. (**F, G**) Representative images (F) and quantification (G) of BrdU foci per PCNA-positive cell in cells transfected with siCTRL or si*REV3L*. (G) shows untreated cells and (F) shows EV cells untreated or fixed 0 or 24 h after cisplatin (2 μM, 24 h), and E6E7 cells untreated. (**I**) Western blot validating REV1 depletion in SCC-1 EV and 16E6E7 cells. α-Tubulin was used as a loading control. Treated conditions are shown in Figure 3. Open circles, CTRL siRNA; filled circles, indicated siRNA. Faded dots, individual cells; colored dots, mean values of at least three biological replicates ± SEM. Scale bars, 5 μm. See also Figure 3.

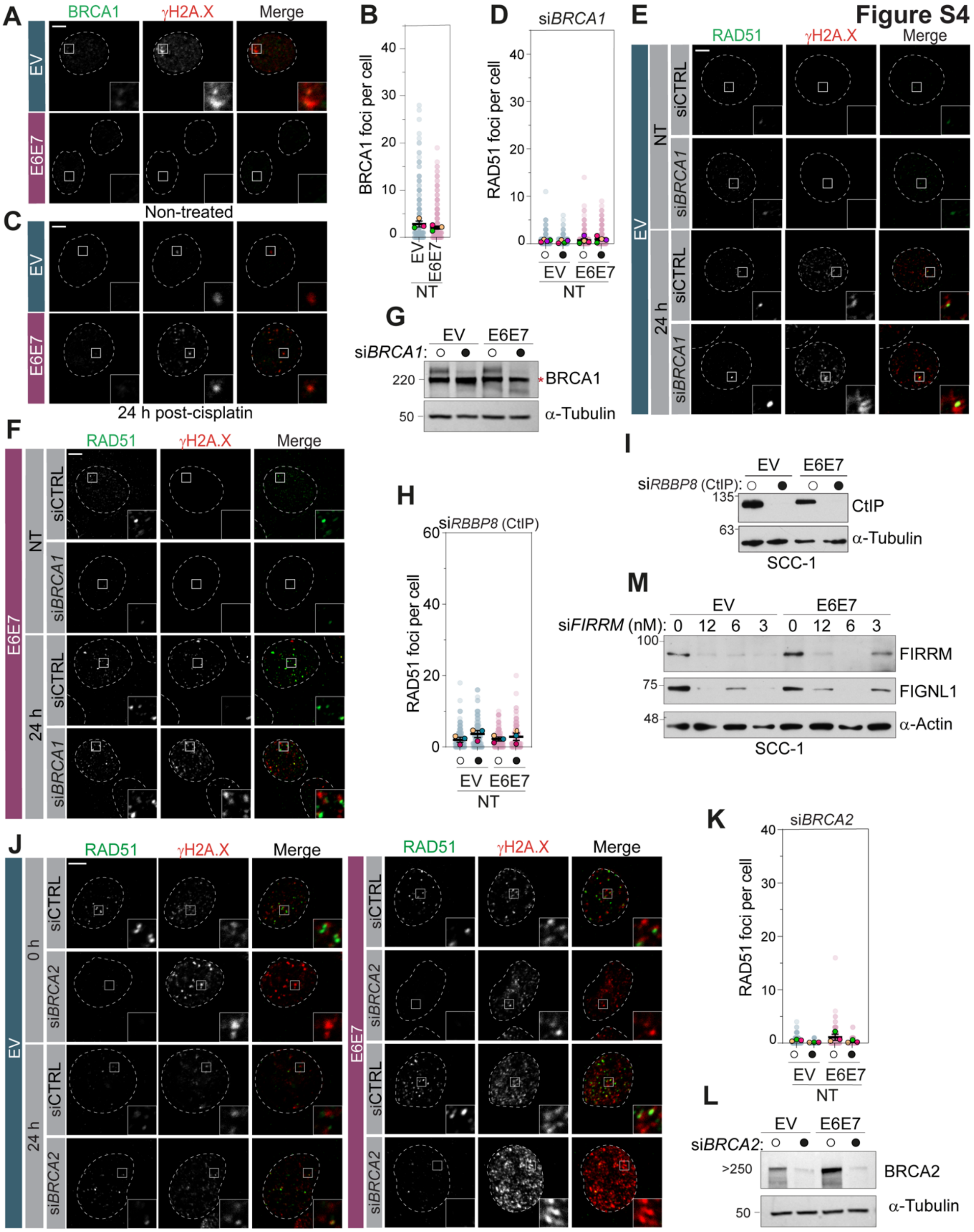


**Figure S4. Validation of depletions by siRNAs and supporting BRCA1 and RAD51 foci data** (**A-C**) Representative images (A, C) and quantification (B) of BRCA1 foci in SCC-1 EV and 16E6E7 cells, untreated (A) or fixed 24 h after cisplatin (250 nM, 24 h) (C). (B) shows untreated cells (NT). (**D-F**) Quantification (D) and representative images of EV (E) and 16E6E7 (F) cells transfected with control (siCTRL) or *BRCA1* siRNA, untreated or fixed 24 h after cisplatin (250 nM, 24 h). (D) shows RAD51 foci in untreated cells. (**G, I, L**) Western blot validating BRCA1 (G), CtIP (I), and BRCA2 (L) depletion in SCC-1 EV and 16E6E7 cells. α-Tubulin was used as a loading control. *, non-specific band. (**H**) RAD51 foci per cell in untreated cells transfected with siCTRL or si*RBBP8*. (**J, K**) Representative images (J) and quantification (K) of RAD51 foci in cells transfected with siCTRL or *BRCA2* siRNA. (J) shows cells fixed 0 or 24 h after cisplatin (250 nM, 24 h), and (K) shows untreated cells. (**M**) Western blot validating FIRRM and FIGNL1 in SCC-1 EV and 16E6E7 cells transfected with the indicated concentrations of si*FIRRM*, showing loss of FIGNL1 upon FIRRM depletion. α-Actin was used as a loading control. Treated conditions are showed in Figure 4. (D, H, K), open circles indicate siCTRL and filled circles the indicated siRNA. Faded dots, individual cells; coloured dots, means of at least three independent biological replicates ± SEM. Scale bars, 5 µm. See also Figure 4.

**Figure S5**

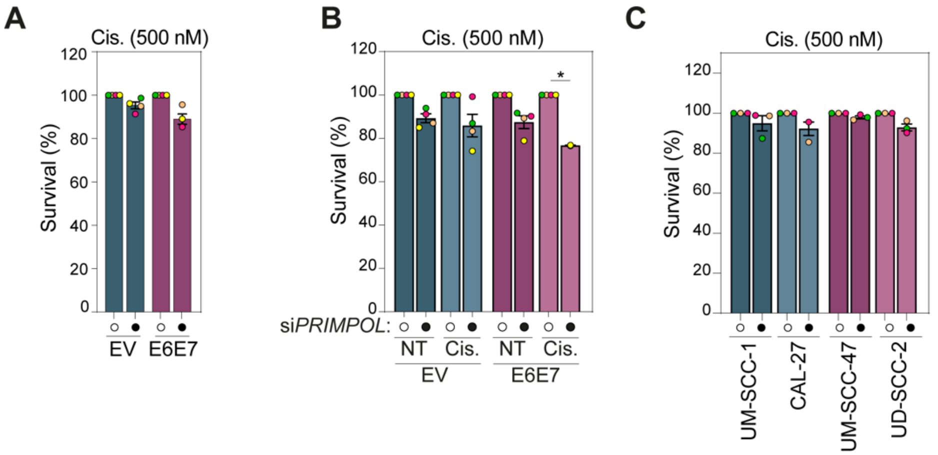


**Figure S5. Cisplatin alone controls in the RAD51 modulation experiments**
(**A**) Survival of SCC-1 EV and 16E6E7 cells untreated (NT, open circle) or exposed to cisplatin (500 nMm 48 h, filled circles), normalized to the untreated condition (100%). (**B**) Survival of the same cells transfected with control (siCTRL, open circle) or *PRIMPOL* siRNA (filled circles), untreated (NT) or exposed to cisplatin (500 nM, 48 h), normalized to the corresponding siCTRL condition (100%). (**C**) Survival of the indicated $HPV^-$ (blue) and $HPV^+$ (pink) cell line untreated (open circles) or exposed to cisplatin (500 nM, 48 h, filled circles), normalized to the untreated condition (100%). Colored dots represent the mean values of at least three biological replicates ± SEM. See also Figure 5.

**Table S1: siRNAs, primers, and gRNAs used in this study**

| siRNA | Sequence (5′–3′) |
|---|---|
| si-FANCA | GGACAGAUCUGCACGGCUCdTdT |
| si-XPF | UGACAAGGGUACUACAUGA-UU |
| si-SLX4 | Dharmacon,SMARTpool<br>UCAAACGGCACUCAGAUAA<br>GCGGAGACUUUGUUGAAAU<br>CAAGUGAGCCCGAGGAACA<br>UCAGAGCCGUCCCAAAUAA |
| si-EXO1 | CAAGCCUAUUCUCGUAUUU |
| si-MUS81 | CAUUAAGUGUGGGCGUCUA |
| si-CTRL OT+ | UGGUUUACAUGUCGACUAAdTdT |
| si-PrimPol | GAGGAAACCGUUGUCCUCAGUGUAU |
| si-Rev3L | Dharmacon,SMARTpool .-6.-( L-006302)<br>J-006302-05: GGAGUUCUCUGCUGAGUUA<br>J-006302-06: ACACAGAAGUUGAGUAAUA<br>J-006302-07: GGAUGUAAGUCCAUGUAUA<br>J-006302-08: GAUAUGAGAUUCAGAUGCA |
| si-REV1 | GAACAGUGACGCAGGAAUAUU |
| si-BRCA2 | GAAGAAUGCAGGUUUAAUAUUdTdT |
| si-BRCA1 | GGAACCUGUCUCCACAAAGdTdT |
| si-RAP80 (UIMC1) | Dharmacon,SMARTpool<br>D-006995-03: AGAGGCAGCUCCUUAAUAA<br>D-006995-04: GCACAAGACUUCAGAUGCA<br>D-006995-20: CCACAAAGAUUGAACGACA<br>D-006995-21: AAGACAAAACAGUCGAAUA |
| si-SMARCAL1 | UUUCACAGAGAAAUGCUUAUGCAGG |
| si-HLTF | GGUGCUUUGGCCUAUAUCAUU |
| si-ZRANB3 | CAAGAGAUAUCAUCGAUUATT |
| si-FIRRM-3 | GAAACGACAACCAGGAUAU |
| si-CtIP | GUACAAGGUUUACAAGUAAdTdT3 |
| si-RAD51 | Dharmacon,SMARTpool<br>D-003530-02: GAAGCUAUGUUCGCCAUUA<br>D-003530-05: GCAGUGAUGUCCUGGAUAA<br>D-003530-07: CCAACGAUGUGAAGAAAUU<br>D-003530-08: AAGCUAUGUUCGCCAUUAA |
| **Primer for RT-qPCR** | **Sequence (5′–3′)** |
| HPV16 E6 Forward | TGTTTCAGGACCCACAGGAGC |
| HPV16 E6 Reverse | CGCAGTAACTGTTGCTTGCAG |
| HPV16 E7 Forward | AGGAGGATGAAATAGATGGTCCAG |
| HPV16 E7 Reverse | CTTTGTACGCACAACCGAAGC |
| HPV33 E6 Forward | AGACACTGAGGAAAAACCACGA |
| HPV33 E6 Reverse | AGGGTTTTTGCATTCCACGC |

| | |
|---|---|
| HPV33 E7 Forward | AGAGGACACAAGCCAACGTT |
| HPV33 E7 Reverse | GTTGTGCTTGTCCATCTGGC |
| TP53 Forward | GCGAGCACTGCCCAACAACA |
| TP53 Reverse | GGATCTGAAGGGTGAAATATTCT |
| BAX Forward | GGTTGTCGCCCTTTTCTA |
| BAX Reverse | CGGAGGAAGTCCAATGTC |
| CDKN1A Forward | TGCCGAAGTCAGTTCCTTGT |
| CDKN1A Reverse | CATGGGTTCTGACGGACATC |
| GAPDH Forward | CAACGTGTCAGTGGTGGACC |
| GAPDH Reverse | TCGTTGAGGGCAATGCCAGC |
| RSP13 Forward | GCTTTACCCTATCGACGCAG |
| RSP13 Reverse | AGGAGTAAGGCCCTTCTTGG |
| Rev3L Forward | TCATGAGAAGGAAAGACACTTTATG |
| Rev3L Reverse | GCTGTAGGAGGTAGGGAATATG |
| PrimPol Forward | TGTGGCTTTGGAGGTTACTGA |
| PrimPol Reverse | TTCTACTGAAGTGCCGATACTGT |
| SLX4 Forward | GATTGACCCTCGCTCCTCTG |
| SLX4 Reverse | CCTTTGGAAAAAGCTAGCGCA |
| PUM1 Forward | TGCGGGAGATTGCTGGACAT |
| PUM1 Reverse | GTGTGGCACGCTCCAGTTTC |
| VAPA Forward | CCGCCCACAGACCTCAAAT |
| VAPA Reverse | CGAGGTGCTGTAGTCTTCACT |
| **gRNA** | **Sequence (5′–3′)** |
| sg-FAM83g (Ctrl) | GAGCCCGGGTCACGCCGCGGT |
| sg-FANCA | GTGGCAAACATAGCTTTCCGA |
| sg-FANCD2 | GCTTAGTAGCCGACTGAAACA |
| sg-FANCI | GAAGCTTGTATTCTGACCCA |
| FAM83g editing validation forward | AACAATAAAGAGGCCCTTTGAGACGG |
| FAM83g editing validation reverse | CTCATCAGGTCTTTCCCGCAGATT |
| **Primer for CRISPR screen** | **Sequence (5′–3′)** |
| TKO library outer-PCR primer FW | AGGGCCTATTTCCCATGATTCCTT |
| TKO library outer-PCR primer RV | TCAAAAAAGCACCGACTCGG |
| TKO library inner-PCR primer trueseq i5 1 | AATGATACGGCGACCACCGAGATCTACACTATAGCCTACACT CTTTCCCTACACGACGCTCTTCCGATCTTGTGGAAAGGACGAG GTACCG |
| TKO library inner-PCR primer trueseq i5 2 | AATGATACGGCGACCACCGAGATCTACACATAGAGGCACACT CTTTCCCTACACGACGCTCTTCCGATCTTGTGGAAAGGACGAG GTACCG |

| | |
|---|---|
| TKO library inner-PCR primer trueseq i5 3 | AATGATACGGCGACCACCGAGATCT ACACCCTATCCTACACT CTTTCCCTACACGACGCTCTTCCGATCTTGTGGAAAGGACGAG GTACCG |
| TKO library inner-PCR primer trueseq i5 4 | AATGATACGGCGACCACCGAGATCTACACGGCTCTGAACACT CTTTCCCTACACGACGCTCTTCCGATCTTGTGGAAAGGACGAG GTACCG |
| TKO library inner-PCR primer trueseq i7 1 | CAAGCAGAAGACGGCATACGAGATCGAGTAATGTGACTGGAG TTCAGACGTGTGCTCTTCCGATCTATTTTAACTTGCTATTTCTA GCTCTAAAAC |
| TKO library inner-PCR primer trueseq i7 2 | CAAGCAGAAGACGGCATACGAGATTCTCCGGAGTGACTGGAG TTCAGACGTGTGCTCTTCCGATCTATTTTAACTTGCTATTTCTA GCTCTAAAAC |
| TKO library inner-PCR primer trueseq i7 5 | CAAGCAGAAGACGGCATACGAGATTTCTGAATGTGACTGGAG TTCAGACGTGTGCTCTTCCGATCTATTTTAACTTGCTATTTCTA GCTCTAAAAC |
| TKO library inner-PCR primer trueseq i7 6 | CAAGCAGAAGACGGCATACGAGATACGAATTCGTGACTGGAG TTCAGACGTGT GCTCTTCCGATCTATTTTAACTTGCTATTTCTA GCTCTAAAAC |
